# Mis-splicing of *Mdm2* leads to Increased P53-Activity and Craniofacial Defects in a MFDM *Eftud2* Mutant Mouse Model

**DOI:** 10.1101/2020.09.22.308205

**Authors:** Marie-Claude Beauchamp, Anissa Djedid, Eric Bareke, Fjodor Merkuri, Rachel Aber, Annie S. Tam, Matthew A. Lines, Kym M. Boycott, Peter C. Stirling, Jennifer L. Fish, Jacek Majewski, Loydie A. Jerome-Majewska

## Abstract

EFTUD2, a GTPase and core component of the splicesome, is mutated in patients with mandibulofacial dysostosis with microcephaly (MFDM). We generated a mutant mouse line with conditional mutation in *Eftud2* and used *Wnt1-Cre2* to delete it in neural crest cells. Homozygous deletion of *Eftud2* leads to neural crest cell death and malformations in the brain and craniofacial region of embryos. RNAseq analysis of embryonic mutant heads revealed a significant increase in exon skipping, in retained introns and enriched levels of *Mdm2* transcripts lacking exon 3. Mutants also had increased nuclear P53, higher expression of P53-target genes, and increased cell death. Their craniofacial development was significantly improved when treated with Pifithrin-α, an inihibitor of P53. We propose that craniofacial defects caused by mutations of *EFTUD2* are a result of mis-splicing of *Mdm2* and P53-associated cell death. Hence, drugs that reduce P53 activity may help prevent craniofacial defects associated with spliceosomopathies.

## Introduction

Splicing is the essential mechanism via which the coding exons of pre-mRNAs are joined and the intervening non-coding sequences are removed. Splicing also contributes to proteomic diversity and regulates mRNA levels via alternative splicing, the process of inclusion and skipping of “alternative” exons (Papasaikas and Valcarcel, 2016; Sperling, 2017; Will and Luhrmann, 2011). The major spliceosome, a ribonucleoprotein complex that regulates and performs the splicing reaction, catalyzes 99% of RNA splicing reactions in human (Wickramasinghe et al., 2015). On the other hand, the minor or U12-dependent spliceosome is only responsible for splicing of approximately 700 minor introns in 666 genes (Olthof et al., 2019).

The major spliceosome is composed of U1, U2, U5, and U4/U6 <u>s</u>mall <u>n</u>uclear ribo<u>n</u>ucleo<u>p</u>roteins (snRNPs) named for their core associated small RNAs (snRNAs). The minor or U12-dependent spliceosome consists of U11, U12, U5, and U4atac/U6atac snRNAs and shares all but 7 proteins with the major spliceosome (Will et al., 2004). The sequential steps in splicing include recognition of a 5’-splice site via the U1/U11 snRNP, binding of U2/U12 snRNP to the branch site on the intron, recruitment of the pre-assembled U4/U6.U5 or U4atac/U6atac/U5 snRNPs, cleavage of the intervening intron, and ligation of the two exons. This process is ubiquitous and yet, mutations in components of the spliceosome can result in tissue-specific abnormalities ranging from retinitis pigmentosa to craniofacial malformations (Beauchamp et al., 2020; Daguenet et al., 2015; Griffin and Saint-Jeannet, 2020; Lehalle et al., 2015).

Mutations in EFTUD2, a GTPase and a core component of the U5 snRNP, are responsible for mandibulofacial dysostosis with microcephaly (MFDM) (OMIM#610536). MFDM patients exhibit a variety of clinical phenotypes of which the most common are: microcephaly, developmental delay, mandibular and malar hypoplasia, as well as external ear anomalies. Amongst the approximately 100 individuals with MFDM reported to date, 86 distinct heterozygous *EFTUD2* mutations have been found, including: deletions/duplications, frameshift, nonsense, splice site, and missense (Czeschik et al., 2013; Deml et al., 2015; Gordon et al., 2012; Huang et al., 2016; Lehalle et al., 2014; Lines et al., 2012; Luquetti et al., 2013; Matsuo et al., 2017; Rengasamy Venugopalan et al., 2017; Sarkar et al., 2015; Smigiel et al., 2015; Voigt et al., 2013; Yu et al., 2018; Zarate et al., 2015). Most of these mutations are *de novo*, although approximately 19% are inherited from an affected parent. Mutations are distributed along the length of the gene, and are presumed to result in a loss of function since no genotype-phenotype correlation is found.

Our aim is to uncover the etiology of abnormal craniofacial malformations in MFDM using mutant mouse models of *Eftud2.* However, despite a 30% reduction of *Eftud2* mRNA and protein levels, mice with constitutive heterozygous deletion of exon 2 of *Eftud2* (*Eftud2^+/-^*) are viable and fertile, and exhibit no craniofacial malformations (Beauchamp et al., 2019). Additionally, RNA sequencing revealed no significant splicing changes in *Eftud2^+/-^* embryos when compared to wild-type littermates. Furthermore, homozygous *Eftud2* (*Eftud2^-/-^*) mutant blastocysts failed to implant, precluding analysis of the role of *Eftud2* during craniofacial development. These and other past studies confirmed that *Eftud2* is an essential gene, but its role in neural crest cells, the precursors of bones and cartilages affected in the head and face of MFDM patients, has not been examined (Beauchamp et al., 2020; Deml et al., 2015; Lei et al., 2017; Wu et al., 2019). Therefore, we have generated a conditional *Eftud2* knock-out mouse (*Eftud2^loxP/+^*) in which exon 2 is flanked by loxP sequences to study the role of *Eftud2* during craniofacial development. In the present study, we evaluated the effect of deletion of exon 2 of *Eftud2* in the developing brain and neural crest cells using the *Wnt1-Cre2* transgenic mouse line. We show that embryos with homozygous mutation of *Eftud2* in the neural crest cells have brain and midface abnormalities and faithfully modeled MFDM. Malformations were associated with alternative splicing of *Mdm2* and increased P53 protein and activity. Knockdown of *Eftud2* increased cell death, and mutant embryos treated with Pifithrin-α, a small molecule inhibitor of P53 (Komarov et al., 1999), showed significant rescue of craniofacial and brain defects. Altogether our data indicate that loss of EFTUD2 in neural crest cells results in differential splicing of *Mdm2* and triggers a P53-associated stress response that leads to craniofacial defects.

## Results

### Homozygous mutation of *Eftud2* in neural crest cells causes craniofacial malformations and embryonic lethality

To characterize the role of *Eftud2* in neural crest cells, we used *Wnt1-Cre2* transgenic mice (Lewis et al., 2013). Since *Eftud2^+/-^* embryos were morphologically and molecularly indistinguishable from wild-type embryos (Beauchamp et al., 2019), matings of *Eftud2^loxP/+^* ; *Wnt1-Cre2^tg/+^* or *Eftud2^+/-^* ; *Wnt1-Cre2^tg/+^* mice to *Eftud2^loxP/loxP^* mice were used to generate homozygous mutant embryos for analysis. From E8.5 to E17.5, embryos of all genotypes were found at the expected Mendelian ratio (Table S1).

Between E8.5 -E10.5, all embryos including homozygous mutants were alive as assessed by the presence of a heart beat. However at E9.5, neural crest cell-specific *Eftud2* homozygous mutant embryos (*Eftud2^loxP/-^* ; *Wnt1-Cre2^tg/+^* or *Eftud2^loxP/loxP^*; *Wnt1-Cre2^tg/+^*) were morphologically abnormal when compared to their control litter mates (*Eftud2^loxP/+^*; *Wnt1-Cre2^tg/+^,* or *Eftud2^loxP/+^*) (Table S2). Homozygous mutant embryos exhibited hypoplasia of the midbrain, the frontonasal prominence, and of the first and second pharyngeal arches when compared to somite-matched controls (Fig. 1a-c).

**Fig. 1.**
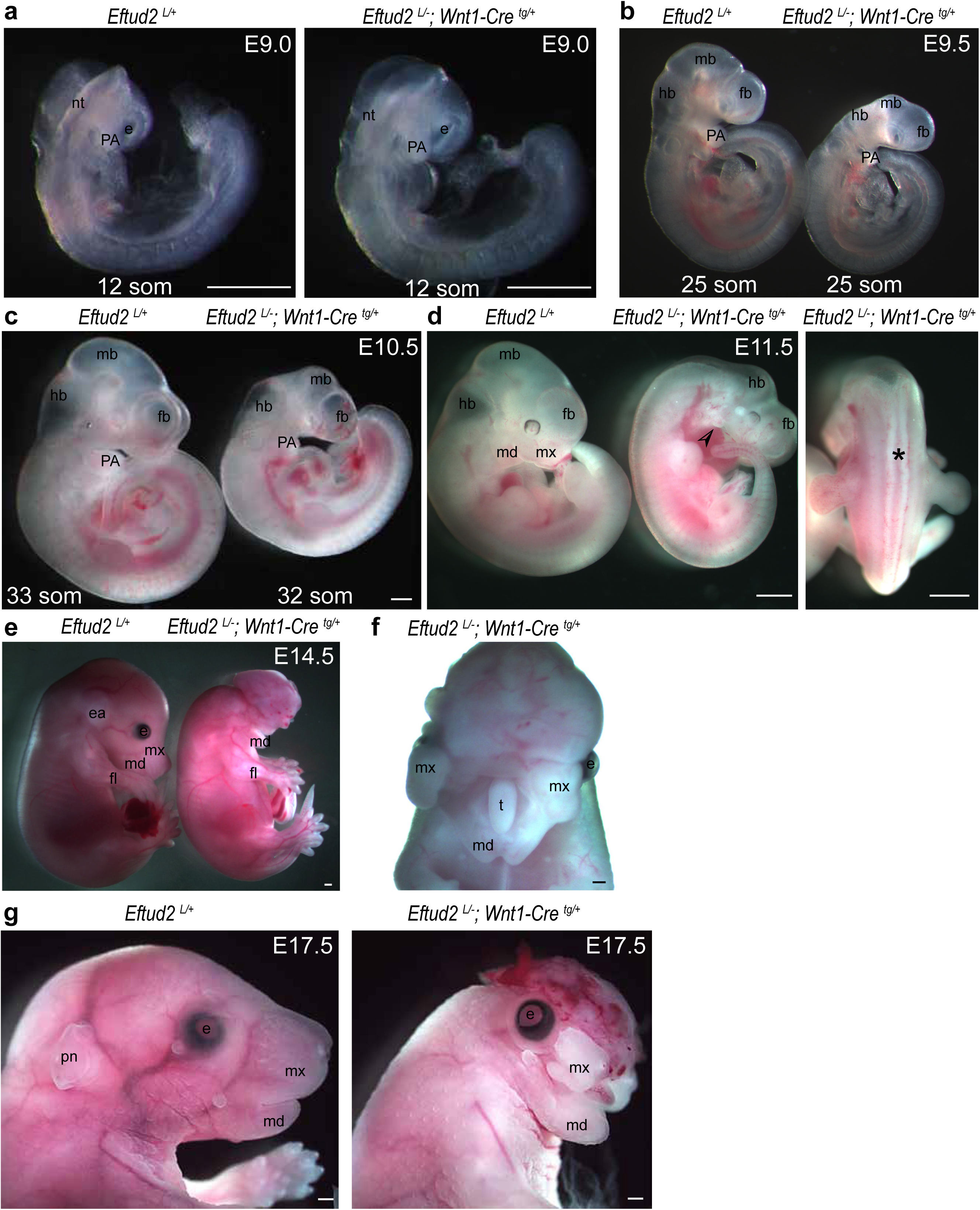
Homozygous mutation of *Eftud2 in* neural crest cells causes craniofacial malformations. **a)** *Eftud2^loxP/-^*;*Wnt1-Cre^tg/+^* mutant embryos are not distinguishable from controls at E9.0. **b)** At E9.5, **c)** E10.5 and **d)** E11.5, *Eftud2^loxP/-^*; *Wnt1-Cre2 ^tg/+^* (*Wnt1-Cre^tg/+^*) mutant embryos exhibit hypoplasia of the midbrain and pharyngeal arches (open arrowhead). The star indicates the open neural tube in an E11.5 mutant embryo. **e)** Embryos found alive at E14.5 had exencephaly and an absence of snout and head structures. **f)** A frontal view of an E14.5 mutant embryo showing a protruding tongue from the oropharyngeal region, a cleft maxillary process and the fused mandibular process. **g)** Image of a dead embryo found at E17.5 showing the hypoplastic lower jaw, cleft maxillary process, missing head structures and absence of eyelid closure. nt: neural tube, PA: pharyngeal arch, hb: hindbrain, mb: midbrain, fb: forebrain, mx: maxillary, md: mandibular prominence, pn: pinnea, fl: forelimb, t: tongue, e: eye. Scale bar=500µm.

Between E11.5 and E17.5, neural crest cell-specific *Eftud2* homozygous mutant embryos showed severe malformations in the head region. At E11.5, 38% of mutant embryos were recovered dead with no heart beat (3/8). The midbrain region was virtually absent in all neural crest cell-specific *Eftud2* homozygous mutants. Brain malformations are most likely a consequence of deletion of *Eftud2* in the midbrain region where *Wnt1-Cre2* is expressed at E8.5 (Lewis et al., 2013). The frontonasal prominence was abnormally shaped, the maxillary and mandibular processes were present but very small (arrowhead), and all had coloboma. Additionally, three of the eight had an open neural tube (star) (Fig.1d).

At E14.5, most neural crest cell-specific *Eftud2* homozygous mutants embryos were dead or resorbed (9/13), and live embryos had exencephaly (n=4/4) (Fig.1e). Among these, eyes could be seen in only one embryo as it was covered by brain tissue in the remaining three. The pinnae or the external ear was not found in any mutant embryos. The frontonasal, maxillary and mandibular regions were hypoplastic and the maxillary process was cleft in all embryos. A hypoplastic tongue was found protruding from the oropharyngeal region (Fig.1f). At E16.5 and E17.5, no neural crest cell-specific *Eftud2* homozygous mutant embryos were found alive (Table S2). In fact, 4/5 were resorbed with little to no embryonic tissue. Only one dead mutant was recovered at E17.5. As shown in Fig.1g, the nasal and the maxillary processes were hypoplastic and much shorter than the mandibular process. The external ear did not form and the skull was almost absent, leaving the brain exposed. Taken together, these data indicate that during craniofacial development, *Eftud2* is essential in neural crest cells not only for differentiation of structures derived from these cells but also for signals that emanate from them.

Death of neural crest cell-specific *Eftud2* homozygous mutant embryos by E14.5 precluded an analysis of craniofacial bone development. Hence, we used Alcian blue staining to examine craniofacial cartilage of E14.5 embryos. This analysis revealed an absence or hypoplasia of cartilages in the head of neural crest cell-specific *Eftud2* homozygous mutant embryos (Fig.2a and Fig. S1). Neural crest derived cartilages which will form the chondrocranium or the base of the skull, and the mid and lower portions of the face, were hypoplastic or missing. Cartilages found in the head of control embryos (stars in Fig.2b) and missing in mutants included: the alas temporalis cartilage (A), the basitrabecular process (B), the orbital cartilage (O), the trabecular cartilage (T), and the paranasal cartilage (PN). Also, one embryo had an absence whereas another one had a reduction of Meckel’s cartilage (M), the precursor of the lower jaw (Fig.2a,b and Fig. S1). In contrast, cartilages in the chondrocranium that were derived from mesoderm were present but reduced in size. Hypoplastic cartilages included: the occipital arch cartilage (OA), the parachordal cartilage (P), and the hypochiasmatic cartilage (Y). The inner ear was also abnormal in both embryos. Identifiable cartilages of the inner ear (Fig.2b) were the cochlear part of the auditory capsule (CO) and the canalicular part of auditory capsule (CA), both of which were abnormally shaped or reduced in size.

**Fig. 2.**
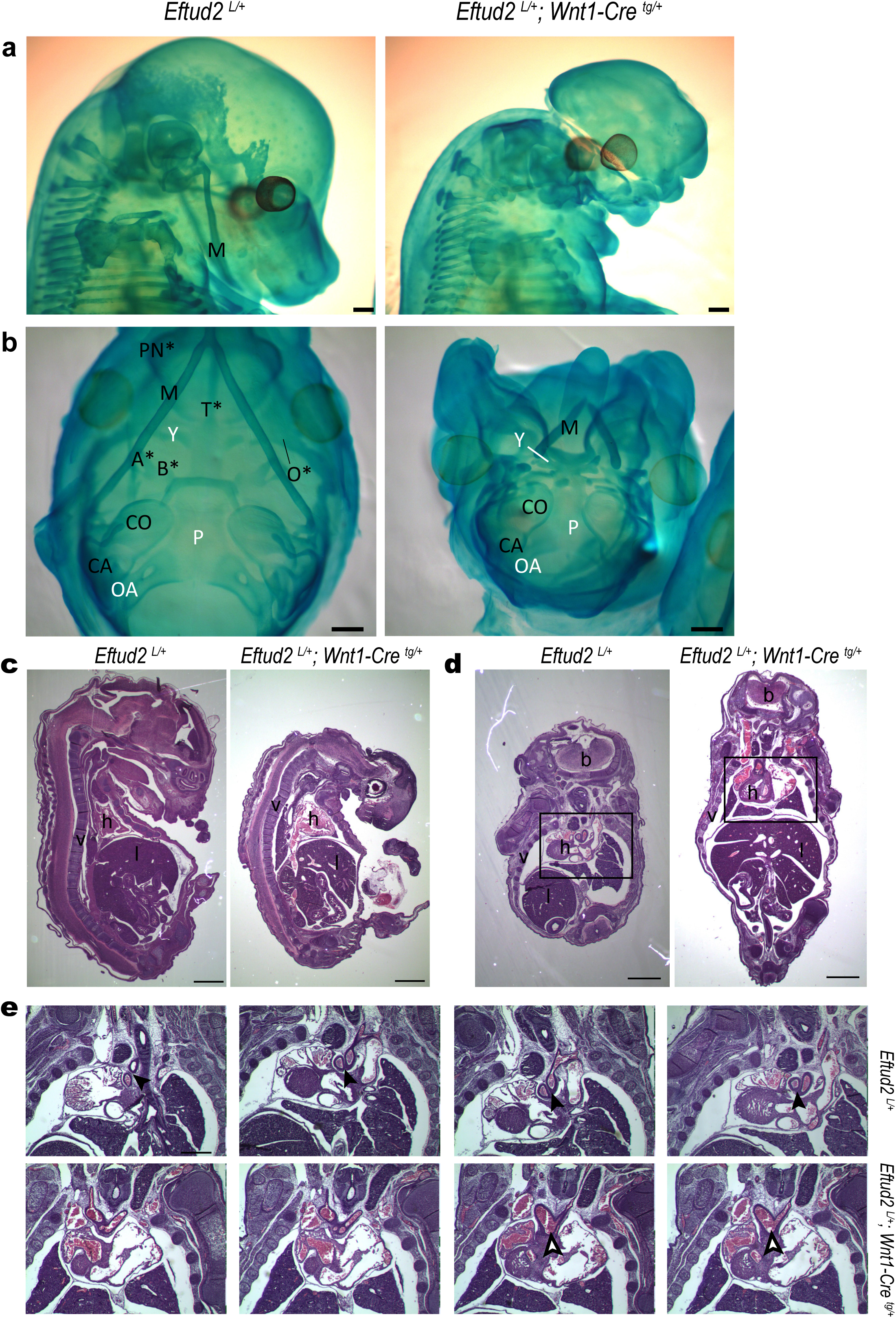
Head and face cartilages are missing or hypolastic in E14.5 *Eftud2^loxP/-^*; *Wnt1-Cre2^tg/+^* mutant embryos. **a)** side view of a control (*Eftud2^loxP/+^*) and *Eftud2^loxP/-^*;*Wnt1-Cre^tg/+^* mutant embryo. Cartilages did not differentiate in the head or face of this mutant embryo. **b)** Ventral view of a control and another *Eftud2^loxP/-^*;*Wnt1-Cre2^tg/+^* mutant embryo showing a hypoplastic Meckel’s cartilage (M) and hypoplasia of the chondrocranium. Structures missing in the mutant are labeled with a star. A: alas temporalis cartilage, B: basitrabecular process, O: orbital cartilage, T: trabecular cartilage, PN: paranasal cartilage, OA : occipital arch cartilage, P: parachordal cartilage, Y: hypochiasmatic cartilage, CO: cochlear part of the auditory capsule, CA: canalicular part of auditory capsule. **c)** Sagittal and **d)** frontal view of H&E stained sections of control and mutant embryos. V: vertebrea, h: heart, l: liver, b: brain. **e)** Higher magnification of serial frontal sections showing a single outflow tract in mutant embryo compared to control (open vs full arrowhead). Scale bar=500µm.

Hematoxilin and eosin staining of the remaining two live embryos confirmed variable levels of hypoplasia and aplasia in the craniofacial region (Fig. 2c, d). In addition, only a single outflow track was found in the heart suggesting that abnormal heart development is responsible for death of neural crest cell-specific *Eftud2* homozygous mutants embryos (Fig. 2e). Aside from this, organogenesis appeared normal in mutant embryos.

### Abnormal trigeminal cranial ganglia formation in neural crest cell-specific *Eftud2* homozygous mutant embryos

In addition to the skeletal elements of the face and head, cranial neural crest cells together with ectodermal placodes, contribute to the formation of the trigeminal (Cranial Nerve; CN V), facial (CN VII), vestibulocochlear (CN VIII), glosopharyngeal (CN IX) and vagus nerves (CN X). To assess the development of the sensory ganglia, we examined expression of the transcription factor *Sox10*, a key regulator of neural crest-derived peripheral glia, at E9.5. At this stage, *Sox10* was expressed in all forming sensory ganglia and in otic vesicles of control embryos (n=3). In neural crest cell-specific *Eftud2* homozygous mutant embryos, *Sox10* expression was reduced in the ganglia, but found in the otic vesicle of all mutant embryos (n=4). Expression in CN V and VII/VIII was variable and found in two and three mutant embryos, respectively. Additionally, *Sox10* expression was found in CN IX in one mutant embryo and not found in CN X or the dorsal root ganglia (stars) of any mutants (Fig.3a).

**Fig. 3.**
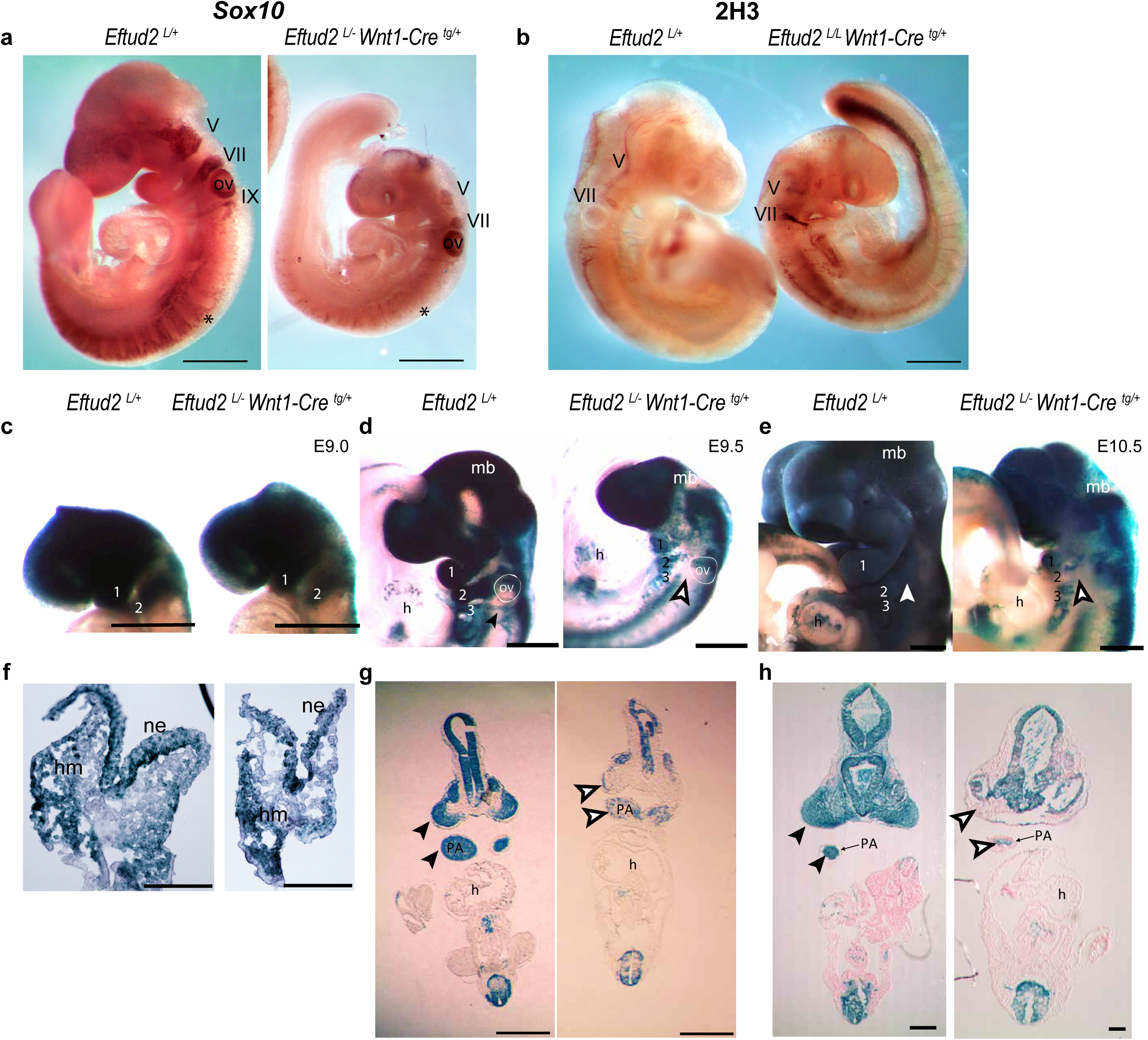
Abnormal trigeminal cranial ganglia formation and reduced neural crest cells in the craniofacial region of *Eftud2^loxP/-^*;*Wnt1- Cre2^tg/+^* or *Eftud2^loxP/loxP^*;*Wnt1-Cre2^tg/+^* mutant embryos. Cranial nerves V and VII can be seen in representative images of control (*Eftud2^loxP/+^*) and mutant E9.5 embryos **a)** after wholemount *in situ* hybridization to detect *Sox10* expression (purple), and **b)** wholemount IHC staining with 2H3 neurofilament antibody (brown). Representative images of control and mutant embryos showing neural crest cells in the developing craniofacial region (blue) **c**) occupy a similar region in E9.0 embryos. **d**) At E9.5 and **e**) E10.5, the region occupied by neural crest cells in the pharyngeal region of *Eftud2^loxP/-^*;*Wnt1-Cre^tg/+^* mutant embryos is reduced. Post-otic streams of neural crest cells destined for pharyngeal arches 3 and 4 (black arrowhead), were absent in *Eftud2^loxP/-^*;*Wnt1-Cre^tg/+^*mutant embryos (white arrowheads). **f-h**) Frontal sections of wholemount stained embryos at **f**) E9.0 **g**) E9.5 and **h**) E10.5. ne: neuroepithelium, hm: head mesenchyme. PA: pharyngeal arch, ov: otic vesicle, h: heart. Star: dorsal root ganglia. Scale bar=500µm.

Similarly, at E9.5 and E10.5, immunohistochemistry using the pan neurofilament antibody 2H3 revealed abnormally formed cranial sensory ganglia in all mutant embryos (n=4). At E9.5, all forming cranio sensory ganglia were distinguishable in controls (n=2) whereas in mutants (n=2), staining was only found in cranial nerve V and cranial nerve VII (Fig. 3b) . At E10.5, CN V and CN VII/VIII were not distinguishable in the mutants, whereas cranial nerve IX and X were stained (n = 2) (Fig S2). Thus, sensory cranial ganglia are formed in neural crest cell-specific *Eftud2* homozygous mutant embryos, however these ganglia are abnormal.

### *Eftud2* mutant neural crest cells migrate to the craniofacial region but fail to survive and/or expand

To determine if neural crest cell generation and/or migration were disrupted by mutation of *Eftud2,* we generated embryos expressing the *R26R* reporter and used X-Gal staining to visualize neural crest cells and their derivatives. At E9.0, the region of X-Gal positive neural crest cells in heads of mutant and control embryos analyzed in wholemounts or after cryosection was comparable (Fig. 3c and f). However, at E9.5 and E10.5, wholemount followed by cryosection revealed that regions of X-Gal positive neural crest cells in the frontonasal region and pharyngeal arches of mutant embryos were significantly reduced when compared to controls (Fig. 3d, e and full vs open arrowheads in g and h). Moreover, the post-otic streams of X-Gal positive neural crest cells from the hindrain to pharyngeal arches 2 and 3 found in control embryos were missing in homozygous mutant embryos (full vs open arrowheads, Fig. 3d and e). Thus, we concluded that either fewer *Eftud2* mutant neural crest cells are generated or fewer of these cells migrate and colonize the developing craniofacial region, either due to a block in survival, expansion and/or differentiation.

### *Eftud2* mutant neural crest cells have increase levels of cell death

To determine if proliferation is perturbed in E9.0 neural crest cell-specific *Eftud2* homozygous mutant embryos, immunofluorescence was used to stain for phospho-histone H3 (PH3) to label cells in mitosis. The number of PH3 positive cells found in mutants (n =4) and controls (n=5) was not significantly different between the two groups (Fig.4a,b). We next examined apoptosis at E9.0 using TUNEL assay to label apoptotic nuclei. Though more TUNEL positive nuclei were found in mutants (n=4) when compared to controls (n=5), this difference was not statistically significant (Fig.4c,d).

**Fig. 4.**
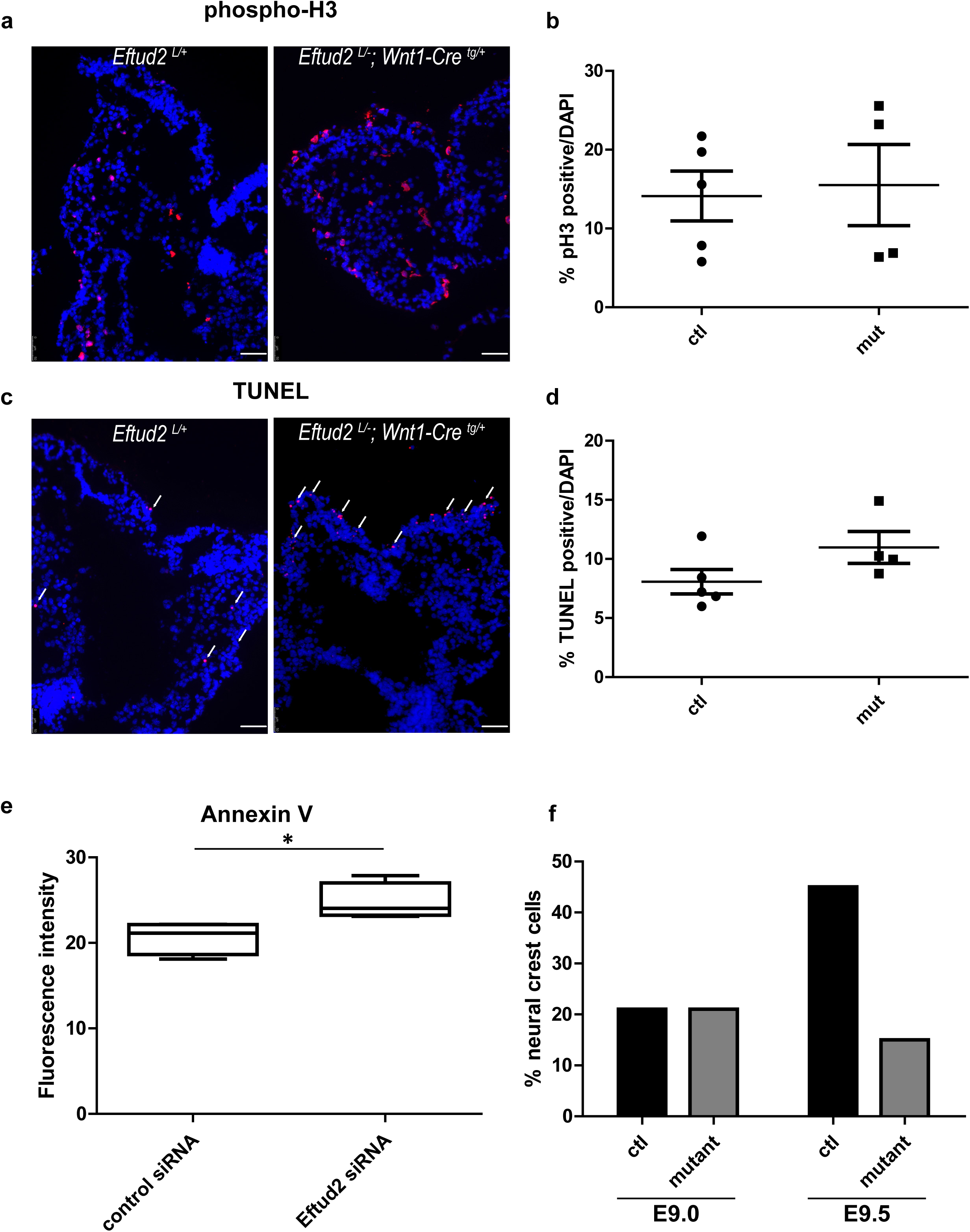
*Eftud2* mutant neural crest cells have increased levels of cell death. **a)** Representative images of E9.0 control (*Eftud2^loxP/+^*) and mutant (*Eftud2^loxP/-^*;*Wnt1-Cre^tg/+^*) embryos showing phospho-histone H3 positive (red) nuclei. **b)** quantification of the % of PH3 stained cells/DAPI. **c)** Representative images of E9.0 control and mutant embryos showing TUNEL positive cells (red) indicated by white arrows. **d)** quantification of the % of TUNEL positive cells/DAPI. **e)** Boxplot of Annexin V fluorescence intensity, in arbitrary units, for control (n=6) and knockdown (n=4) 09-1 neural crest cells 24-30 hours post-transfection with negative control or *Eftud2* siRNAs, respectively. A Kruskal-Wallis rank sum test was used to determine the significance associated with the difference between the treatment groups (* p≤0.05). **f)** The percentage of neural crest cells in heads of control and mutant embryos at E9.0 and E9.5 using the exclusion of exon 2 of *Eftud2* from the RNAseq data as a surrogate for the neural crest cells population (see Methods).

To further test if reducing levels of *Eftud2* increase apoptosis of neural crest cells, O9-1 cells transfected with *Eftud2* siRNA or control siRNA were stained with Annexin V. A significant increase in Annexin V staining intensity was found in cells with knockdown of *Eftud2,* when compared to controls. These data indicate that neural crest cells with reduced levels of *Eftud2* undergo apoptosis (Fig.4e).

### Altered transcriptional landscape in *Eftud2* homozygous neural crest mutants

EFTUD2 is a core component of the spliceosome so we next sought to investigate whether its absence results in detectable defects or alterations in pre-mRNA splicing. We used ribo-depleted RNA isolated from the craniofacial regions of E9.0 and E9.5 neural crest cell-specific *Eftud2* homozygous mutant embryos (*Eftud2 ^loxP/-;^Wnt1-Cre2 ^tg/+^*) and controls (*Eftud2 ^loxP/+;^; Wnt1-Cre2 ^tg/+^*) for RNAseq (see Methods). We then used the rMATS computational package to identify splicing differences between *Eftud2* homozygous mutants and controls, including skipped exons, retained introns, the use of alternative 5’ or 3’ splice sites, and mutually exclusive exons. We expected the differences that most directly reflect the effects of *Eftud2* deletion to be present at E9.0 (before the onset of first phenotypic differences), whereas differences at E9.5 may reflect a combination of both direct and secondary effects resulting from altered development.

Exon skipping was the most frequent type of differential alternative splicing event with 567 (E9.0) and 380 (E9.5) differential exon skipping events (FDR < 0.05) (Fig.5a and Fig. S3a). The most significant detected event was the skipping of exon 2 of *Eftud2;* while this is not an actual alternative splicing event, but the result of the targeted deletion, it serves as a convenient positive control indicating the validity of the analysis. Exclusion of exon 2 of *Eftud2* from the RNA sequencing data also allows us to estimate the fraction of neural crest cells present within the mutant and controls craniofacial regions (see Methods) to be 21% at E9.0. At E9.5, this proportion increases to 45% in the control embryos, but falls to 15% in the mutants, providing independent confirmation for the inability of *Eftud2* mutant neural crest cells to survive and expand (Fig.4f). Among legitimate skipped exon events, there was a clear and significant tendency for increased exon skipping in mutants at both stages (Fig. 5a and S3a). No difference was found for mutually exclusive exons (Data not shown). Among the remaining alternative splicing differences, there was a slight significant preference for the usage of upstream splice sites (alternative 5’ and 3’ sites) at E9.0, however this trend was not observed at E9.5 (Fig. 5c, d and Fig. S3c, d). Intron inclusion was preferentially increased in the controls, relative to the mutants, at both stages, but notably the overall number of detected retained introns was small (Fig.5b and S3b).

**Fig. 5.**
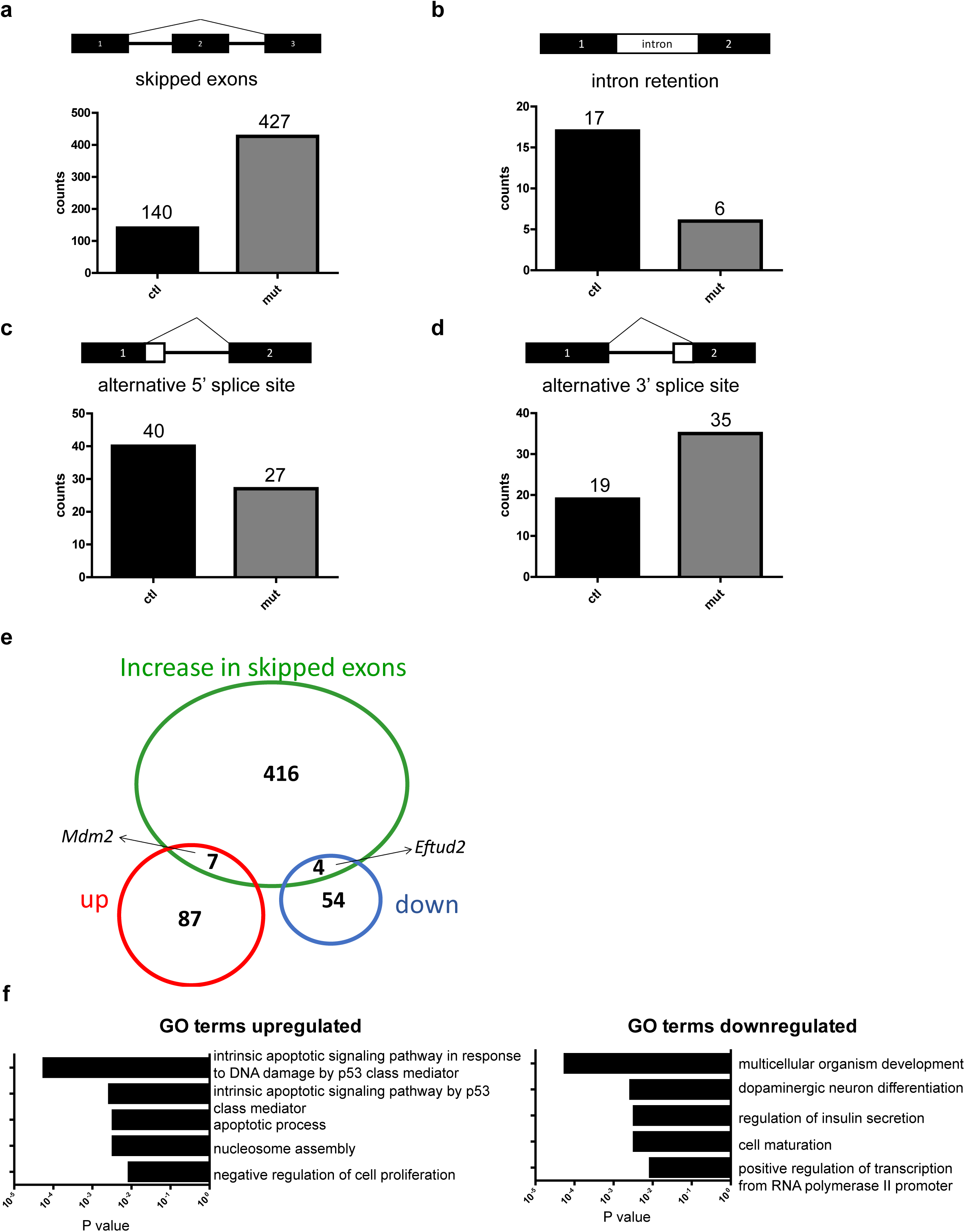
RNAseq analysis at E9.0 reveals increased exon skipping and upregulation of the P53 pathway. The number of various alternative splicing events is represented and a sign test performed to compare the tendency for each type of event to occur preferentially in mutants versus controls **a**) skipped exons (P<0.0001) **b**) intron retention (P<0.05, more frequent in controls) **c**) alternative 5’ splice site **d**) alternative 3’ splice site (P<0.05 when jointly testing 5’ and 3’ splice site preference for the usage of downstream site). **e**) Venn Diagram showing genes with skipped exons in mutant embryos that were also upregulated (including *Mdm2*) or downregulated (including *Eftud2*) **f**) GO terms analysis of genes that were upregulated or downregulated in mutant vs controls.

Retained introns are traditionally difficult to identify, and to further explore intron retention differences, we used IRFinder (Middleton et al., 2017), an approach specifically designed for this purpose. The most significant differential intron retention events, at both developmental stages, were the 2 introns flanking exon 2 of *Eftud2*; again, although this is not a true splicing difference, it is a positive control validating the approach. We used a nominal p-value < 0.05 as a selection criterion in order to study any tendencies for intron inclusion that may result from *Eftud2* deletion. At stage E9.0, we detected more retained introns in the mutant (436 out of 550, p = 2.4e-45). Paradoxically, this tendency was reversed at E9.5 with significantly lower numbers of retained introns in the mutant (238 out of 769, p = 1.53e-26).

To further investigate possible mechanisms through which EFTUD2 may affect splicing, we focused on the most consistent and abundant trend in exon-skipping in the mutants and considered the E9.0 stage, where the contribution of neural crest cells to the head region was comparable in mutants and controls. We compared splice site strength and nucleotide profiles between exons that were preferentially skipped in the mutant (representing the types of splice sites that may be less efficiently recognized in the absence of ETUD2) to the splice sites that were preferentially included in the mutant (which we believe would represent secondary downstream splicing differences, likely independent of EFTUD2). However, we did not observe any difference in splice sites characteristics (Fig. S4), suggesting that EFTUD2 may not be involved in the process of splice site recognition. To explore this idea further, we asked whether the absence of EFTUD2 may result in more frequent usage of unannotated or cryptic splice sites. Again, we did not detect any difference in unannotated spliced-site usage, suggesting that the absence of EFTUD2 does not result in mis-directing the splicing machinery to atypical splicing signals (Fig. S5).

Alternative splicing events could potentially lead to changes in gene expression with the introduction of exons containing premature termination codons (PTC). Therefore, we compared log fold change of transcripts with skipped exons or retained introns. No significant difference was found in the expression level of these genes (Fig. S6). Next, we specifically identifed those skipped exons events that were predicted to introduce PTCs and asked whether increased PTC corresponded to decreased expression of host genes. Among the 82 PTC-causing exons that were significantly more included in mutants than controls, 48 were associated with increased, while 34 were associated with decreased gene expression (Fig. S6). Furthermore, the magnitude of gene expression differences was not different between the genes with increased and decreased expression. Similar analysis of events with retained introns suggest that in heads of neural crest cell-specific *Eftud2* homozygous mutant embryos, the alternative splicing events found did not correlate with a significant overall change in gene expression. Thus, our results do not support the role of mutant EFTUD2-associated splicing defects in downregulation of genes expression.

To determine if splicing is also altered in patient derived cells, we compared results of RNAseq analysis of lymphoblastoid B-cell lines (LCL) from four unrelated MFDM patients to one control. We identified 35 common splicing changes in these cell lines that were not seen in the control (3 3’SS; 6 5’SS; 21 SE; 5 MXE) (Table S3;S4). While this analysis was underpowered and the transcriptome of immortalized LCLs is not likely relevant models to human neural crest cells in MFDM, the existence of recurrent splicing changes in these human models supports a mechanism of MFDM involving splicing changes.

### Genes in the P53 – pathway are upregulated in *Eftud2* mutant neural crest cells

We next analyzed our data set to identify genes upregulated and downregulated in the heads of neural crest cell-specific *Eftud2* homozygous mutant embryos. At E9.0, 94 transcripts were upregulated and 58 transcripts were downregulated in mutants when compared to controls (Fig.5e). An analysis of gene ontology terms significantly associated with these transcripts revealed the two most significant upregulated pathways to be intrinsic apoptotic signalling pathway in response to DNA damage by P53 class mediator and intrinsic apoptotic signalling pathway by P53 class mediator (Fig.5f). The two most significant downregulated pathways were multicellular organism development and dopaminergic neuron differentiation (Fig.5f). A comparable analysis using KEGG revealed P53 signalling to be the most significantly upregulated pathway (data not shown). Similarly, the P53-signalling pathway was among the the top pathways significantly upregulated in LCLs from MFDM patients (Table S5).

Using RT-qPCR, we confirmed significant increases of P53 – pathway genes: *Ccng1*, *Phlda3*, *Trp53inp1* and *Mdm2*, in the head of neural crest cell-specific *Eftud2* homozygous mutant embryos (Fig.6a). To test if reduced levels of *Eftud2* in neural crest cells also led to increased levels of P53 target genes, we transfected mouse neural crest cells from line O9-1with an *Eftud2* siRNA to reduce levels of this gene. RT-qPCR revealed that expression of *Ccng1*, *Phlda3*, and *Trp53inp1* were also significantly increased in cells with *Eftud2* knockdown when compared to controls (Fig. S7a). These data indicate that reducing levels of *Eftud2* results in increased P53 activity.

### Mutations of *Eftud2* lead to increase skipping of exon 3 of *Mdm2,* a master regulator of P53, and an increase in nuclear P53

Though a global change in gene expression was not associated with the alternative splicing events found in neural crest cell-specific *Eftud2* homozygous mutant embryos, our data confirmed increased skipping of the deleted exon 2 in *Eftud2* which was associated with dowregulated expression of this gene (Fig.5e). The list of genes with increased alternative splicing also included increased skipping of exon 3 of *Mdm2*, the master regulator of P53 (Fig.5e). Alternative splicing of exon 3 of *Mdm2* produces a shorter protein that can no longer bind to MDM4 and leads to stabilization and upregulation of P53 (Giglio et al., 2010). Therefore, we used RT-PCR to examine alternative splicing of this gene in E9.0 embryos. Using primers targeting exon 2 and 6 of *Mdm2*, the proportion of *Mdm2* transcripts lacking exon 3 was found to be significantly increased in *Eftud2* homozygous neural crest mutant embryos when compared to controls (Fig.6b). Similarly, O9-1 neural crest cells with knockdown of *Eftud2* (Fig.6c) and patient LCLs (Fig.6d) also showed a significant increase in *Mdm2* transcripts lacking exon 3 when compared to controls. To assess if the increased splicing of *Mdm2* exon 3 was responsible for the overactivation of the p53 pathway in neural crest cells, we overexpressed full-length *Mdm2* in cells transfected with *Eftud2* siRNA and evaluated splicing of *Mdm2* and expression of *Ccng1*, *Trp53inp1* and *Phlad3*. No difference was found in splicing of *Mdm2* when full-length *Mdm2* was overexpressed (Fig. S7b). But, as shown in Fig.6e, although not significant, there was a tendency towards decreased expression of all genes in the p53 pathway in cells with *Mdm2* overexpression.

**Fig. 6.**
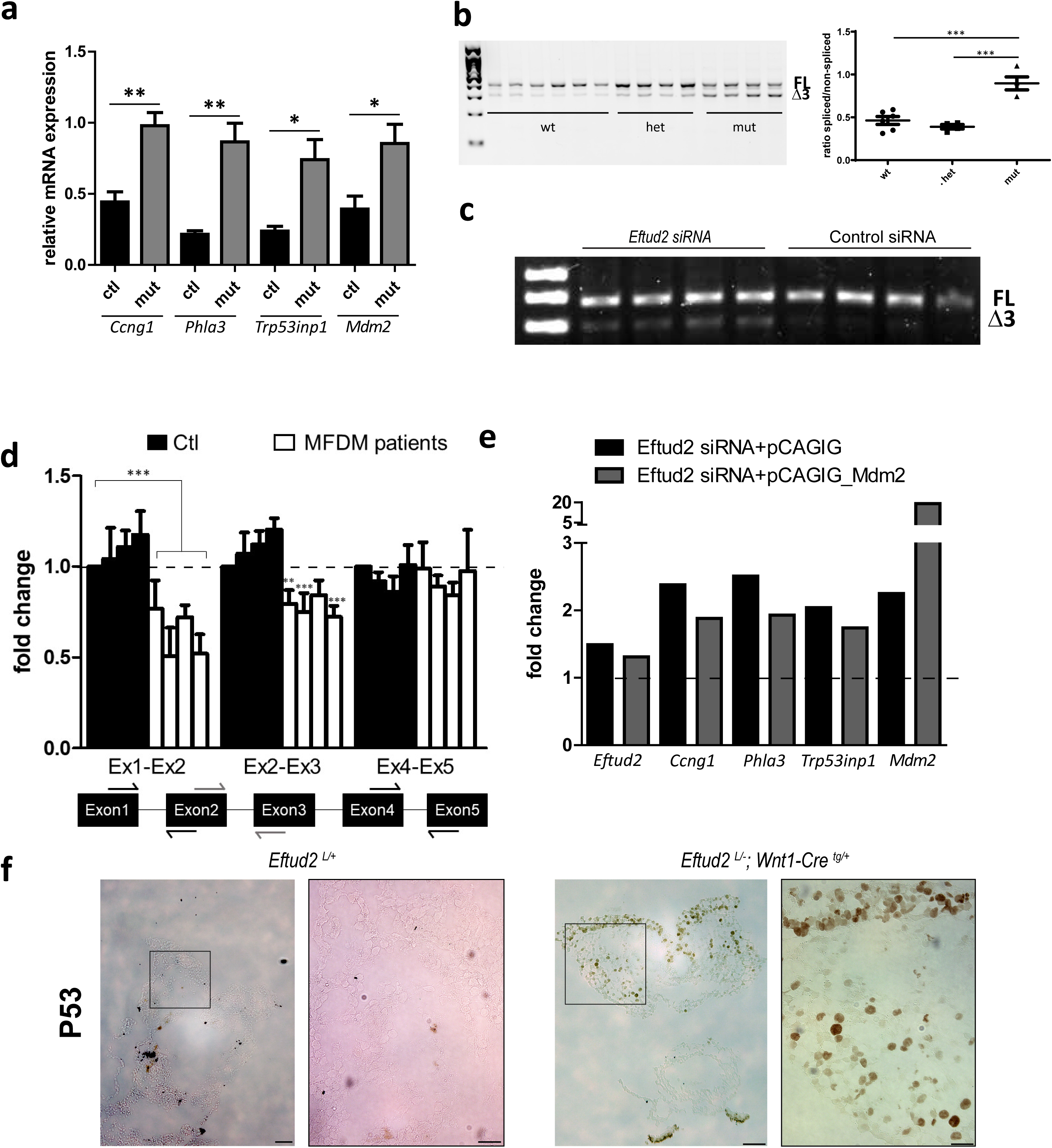
Alternatively spliced and overexpressed *Mdm2* leads to increased p53 when *Eftud2* is mutated. **a)** RT-qPCR showing increased expression of P53 target genes in E9.0 *Eftud2^loxP/-^*;*Wnt1-cre^tg/+^* mutant embryos. **b)** RT-PCR using primers flanking exon 3 of *Mdm2* confirms a significant increase in transcripts with skipped *Mdm2* exon 3 in E9.0 *Eftud2^loxP/-^*;*Wnt1-cre^tg/+^* mutant embryos. Left, representative gel image and right, quantification of splicing of *Mdm2* exon 3. FL: full-length transcript Δ3: transcript without exon 3 **c)** Representative RT-PCR using the same primers for *Mdm2* in O9-1 cells showing increase of the *Mdm2* transcript with skipped exon 3 in cells transfected with *Eftud2* siRNA compared to control siRNA. **d)** RT-qPCR showing fold change of *Mdm2* transcripts with exons 1 – 4 in LCL lines isolated from MFDM patients when compared to controls. Schematic highlights primer flanking locations. p-values calculated from ΔΔCt levels (ANOVA). Mean values with S.E.M. error bars are shown, n = 3. **e)** Fold change in *Eftud2*, *Ccng1*, *Trp53inp1*, *Phlda3*, and *Mdm2* levels in O9-1 neural crest cells transfected with *Eftud2* siRNA and pCAGIG (black bars) or pCAGIG_*Mdm2* (grey bars) expression vectors. Dashed horizontal lines denote expression in control cells transfected with pCAGIG. Data presented as average fold-change. **f**) Representative image of sections of E9.0 control (*Eftud2^loxP/+^*) and *Eftud2^loxP/-^*;*Wnt1-cre^tg/+^* mutant embryos showing increased P53 (brown) positive cells after IHC. Scale bars, lower magnification=75μm, higher magnification=25µm. *P<0.05 **P<0.01, ***P<0.001.

Increased skipping of exon 3 of *Mdm2* was also previously shown to lead to increased nuclear P53 (Van Alstyne et al., 2018). To determine if this is the case in mutant embryos, we performed immunohistochemistry to detect P53 in E9.0 mutant (n=4) and control (n=5) embryos. P53 was barely detectable in control embryos, while striking expression of nuclear P53 was found in the neuroepithelium and surrounding head mesenchyme of all E9.0 neural crest cell-specific *Eftud2* homozygous mutant embryos (Fig.6f). These findings indicate that reduced levels of *Eftud2* leads to alternative splicing of *Mdm2*, and that this in turn results in increased accumulation of nuclear P53 and expression of P53 target genes.

### Decreasing P53 activity with Pifithrin-⍺ rescues a subset of craniofacial defects in *Eftud2* neural crest mutants

Our data show that neural crest cells with mutation of *Eftud2* have increased P53 activity and consequently undergo cell death. Therefore, we postulated that reducing P53 activity could protect these cells from death and rescue morphological abnormalities found in neural crest cell-specific *Eftud2* homozygous mutant embryos. To test if these defects could be prevented by reducing P53, we treated pregnant females with 2.2mg/mg pifithrin-α or DMSO/PBS (vehicle) daily through intra-peritoneal injection, starting at E6.5 until E8.5 and collected embryos at E9.5. Pifithrin-⍺ is a reversible chemical inhibitor of P53 activity that is thought to protect cells from P53-induced cell death by decreasing nuclear P53 activity or modulating its nuclear import/export (Komarov et al., 1999). We first evaluated midbrain development. As shown in Fig.7a, control embryos treated with vehicle (n=8/8) or pifithrin-α (n=12/12) showed normal midbrain development (highlighted in green and data not shown). On the other hand, a subset of neural crest cell-specific *Eftud2* homozygous mutant embryos treated with pifithrin-α showed a significant improvement in midbrain development (n= 3/11) when compared to vehicle and untreated mutant embryos (n= 0/41) (p=0.0139, Fisher exact test). We next evaluated the perimeter of the first pharyngeal arch (black dotted lines) of the same groups of embryos (Fig.7a). Treatment with pifithrin-α did not impact the perimeter of pharyngeal arch 1 of control embryos, however, pifithrin-α treatment significantly increased the size of the first pharyngeal arch in neural crest cell-specific *Eftud2* homozygous mutant embryos, p<0.05 by t-test (Fig.7b). Taken together, these data show for the first time that reducing P53 in a mouse model carrying the *Eftud2* mutation can, at least partly, prevent craniofacial abnormalities.

**Fig. 7.**
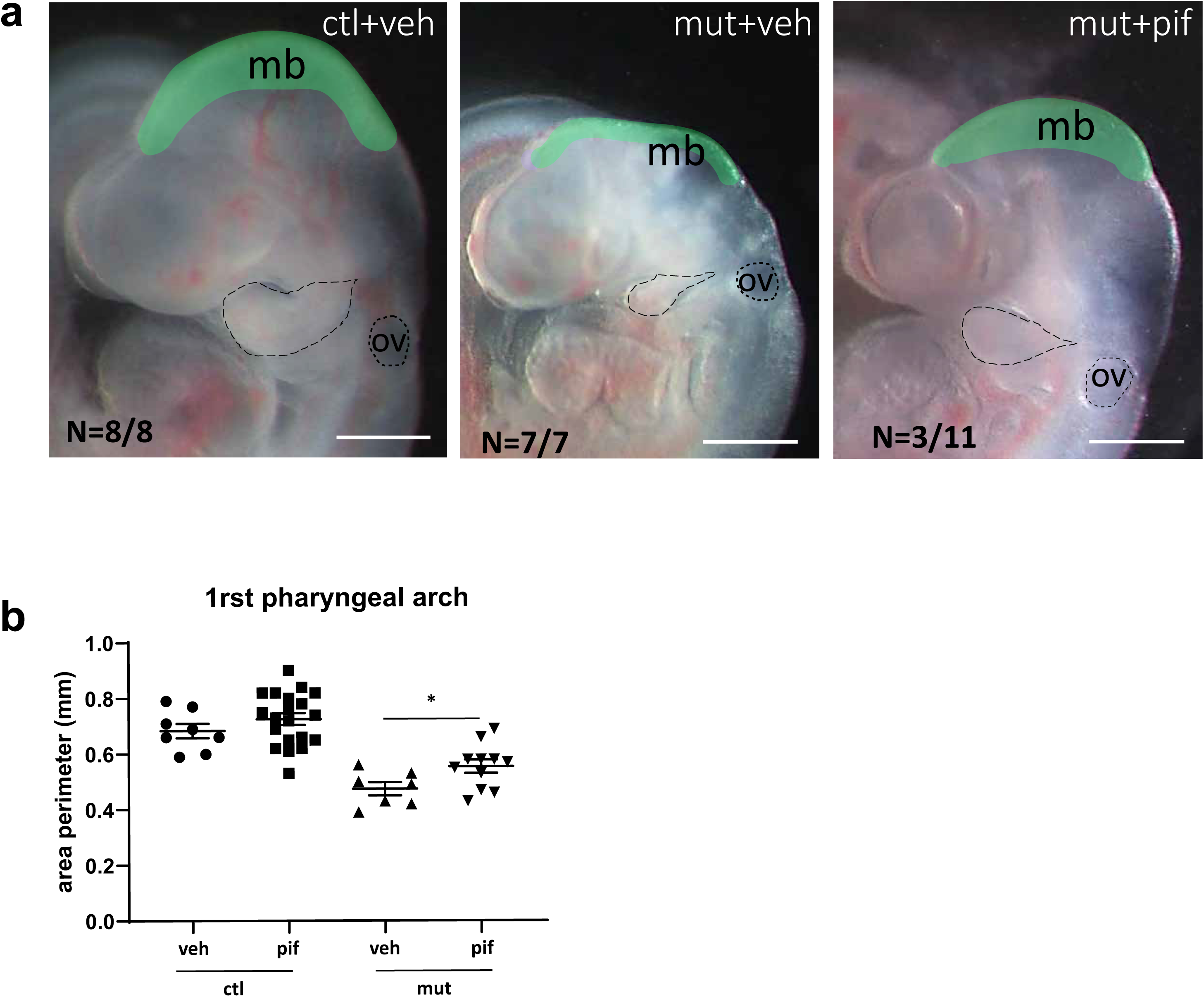
Pifithrin-α partially rescues craniofacial abnormalities in *Eftud2^loxP/-^*;*Wnt1-cre^tg/+^* mutant embryos. Pregnant females were treated daily with vehicle (veh:2% DMSO/PBS; n=3) or 2.2mg/kg pifithrin-α (pif) (n=7) I.P. for 3 days from E6.5 to E8.5, and embryos were collected at E9.5. **a**) Representative pictures of control embryos (*Eftud2^loxP/+^* and *Eftud2^loxP/+^*; *Wnt1-cre^tg/+^*) treated with vehicle (ctl+veh), neural crest mutant embryos (*Eftud2^loxP/-^*;*Wnt1-cre^tg/+^*) treated with vehicle (mut+veh), and neural crest mutant embryos treated with pifithrin-α (mut+pif). The latest were categorized by their head morphology (green area) as normal, n=3/11 or abnormal, n=8/11. **b**) Perimeter of first left pharyngeal arches (black dotted lines). *P<0.05 by t-test. Scale bar=500µm.

## DISCUSSION

Haploinsufficiency of *EFTUD2* was first reported in 2012 to be the cause of MFDM (Lines et al., 2012). Patients with MFDM exhibit a variety of clinical findings and the most common are craniofacial defects such as microcephaly, developmental delay, mandibular and malar hypoplasia as well as external ear anomalies. In addition, as no correlation is found between patient’s genotypes and their clinical characteristics, these mutations are predicted to be loss-of-function mutations and suggest that the developing craniofacial region is more sensitive to reduced levels of EFTUD2 when compared to other embryonic cell types (reviewed in (Beauchamp et al., 2020)). Here, we report that altered splicing caused by the loss of *Eftud2* results in the reduced inclusion of exon 3 of *Mdm2* and nuclear accumulation of P53 in the neural crest cells. This triggers cell death, decreasing the number of neural crest cells that can contribute to craniofacial development. Our findings provide a mechanistic link between EFTUD2 splicing and MFDM. We hypothesize that variability in the number of surviving neural crest cells is responsible for the range of clinical findings reported in MFDM patients.

### Mutations of *Eftud2* in neural crest cells model MFDM

A number of animal models carrying *Eftud2* mutations have been reported in zebrafish and in mouse (Beauchamp et al., 2019; Deml et al., 2015; Lei et al., 2017; Wu et al., 2019). Three zebrafish models targeted *eftud2* gene in different ways: one TALEN-derived deletion in the 5^th^ exon, c.378_385, disrupts part of the GTP binding domain (Deml et al., 2015), a 5 bp deletion in exon 2 also generated using TALEN mutates part of the N-acidic terminal domain (Wu et al., 2019), and an N-ethyl-N-nitrosourea (ENU) induced nonsense C-T mutation in exon 24 of *eftud2* mutates part of the conserved EF 2 domain IV (Lei et al., 2017). These mutants were all predicted or shown to have a decrease or absence of eftud2 protein. Interestingly, all the homozygous mutant zebrafish embryos exhibited similar phenotypes: smaller head size, smaller eyes, curved body tail, and embryonic lethality while heterozygous mutants were normal. A shortened jawbone and deformity of Meckel’s cartilage was found in heterozygous fish with mutation of the N-acidic terminal domain (Wu et al., 2019).

We reported a unique mouse model with heterozygous *Eftud2* mutation in which exon 2, part of the N-terminal acidic region, was deleted. In an outbred genetic background, heterozygous mice were viable and fertile and showed no major phenotypic abnormalities whereas homozygous mutants arrested pre-implantation (Beauchamp et al., 2019). However, although heterozygous pups on the inbred C57Bl/6 genetic background showed no obvious craniofacial malformations, a small but significant number of these pups were lost between birth and weaning. The mouse model described herein is the first to show craniofacial defects similar to those found in patients with mutations in *EFTUD2*, suggesting that the cell of origin of *Eftud2*-associated craniofacial defects is the neural crest cells. Neural crest cells contribute extensively to the formation of the bones of the face and skull (Gilbert, 2010), forming mostly the anterior part of the skull (McBratney-Owen et al., 2008).

### Splicing defects in *Eftud2* mutant models are associated with increased P53 activity

EFTUD2 is a core component of the U5 spliceosome complex that binds to the 5’ (upstream) and 3’ (downstream) exons during the two splicing steps. U5 binds the 5’ exons intermediate generated after the first splicing reaction to tether and align it to the 3’ exons for the second step of the splicing reaction (Newman, 1997). GTP-bound EFTUD2 is predicted to activate the RNA helicase involved in the first step of splicing (Brenner and Guthrie, 2006) and formation of the lariat intron intermediate. Additionally, dephosphorylation of EFTUD2 is predicted to contribute to spliceosome dissassembly (Agafonov et al., 2016; Frazer et al., 2008). However, recent crystal structures of the pre and post-catalytic spliceosome suggests that EFTUD2 interaction with PRP8 may predominantly serve as a “relay station” which enables efficient splicing (Jia et al., 2020).

RNA sequencing analysis of zebrafish with a mutation in the 5^th^ exon of eftud2 revealed a transcriptome-wide RNA splicing deficiency. These mutants had a large number of transcripts with increased intron-retention and exon-skipping, leading to overwhelming nonsense-mediated RNA decay (Lei et al., 2017). The increased stress on the non-sense mediated pathway was predicted to activate P53 and result in apoptosis (Lei et al., 2017). Although a partial rescue of mutant phenotypes and apoptosis was observed when a key gene in the nonsense-mediated pathway, *upf1*, was overexpressed (Lei et al., 2017) or when p53 was inhibited by morpholinos in two different mutant models (Lei et al., 2017; Wu et al., 2019), no direct link was made between splicing and P53 activation.

Recently, Wood *et al*. (Wood et al., 2019) generated a loss-of-function heterozygous mutation in *EFTUD2* in the human embryonic kidney cell line HEK293K. Reduction of EFTUD2 protein caused a decrease in proliferation and increased sensitivity to endoplasmic reticulum (ER) stress. Interestingly, RNA sequencing analysis revealed widespread changes in expression of a number of genes, as well as general splicing differences, most of which were skipped exons. They proposed that the observed splicing differences may indicate “mis-splicing” which could lead to an overwhelming burden of unfolded proteins in the ER, ultimately inducing apoptosis. Similarly, experiments aimed at evaluating the role of *EFTUD2* on human osteoblast proliferation showed that *EFTUD2* knockdown in HCO (human calvarial osteoblast) and in HC-a (human articular chondrocyte) cells reduced their proliferation and differentiation (Wu et al., 2019). Again, RNA sequencing uncovered that the P53 pathway was the most severly affected.

Our study also shows that there is a significant increase in exon skipping in heads of *Eftud2* neural crest cell mutant embryos as compared to controls. However, other alternative splicing events are more difficult to interpret. At E9.0 we do observe a slight tendency towards higher intron retention in the mutants, which is consistent with observations made in other model systems; however, this trend is not seen at E9.5. Other events, such as alternative 5’ and 3’ splice site usage and mutualy exclusive exons also do not show consistent trends across stages. It is likely that our system has limited power to detect the less frequent types of splicing defects, since we estimate that mutant neural crest cell constitute only a fraction of the tissue profiled by RNA-seq. Focusing on the most consistent and abundant category - exon skipping events - we asked whether there is any tendency towards inefficient recognition of weak splice sites in the mutants, but we did not observe any difference between splice site strength of exons with decreased, as compared to increased inclusion. This observation is consistent with the function of EFTDU2 as a GTP-ase and its involvement in maintaining splicing efficiency, but not the recognition of the splice site signals. Contrary to previous hypotheses, we did not find global disruption in expression of genes that contained differential splicing events predicted to lead to premature termination and subject to nonsense-mediated decay. Furthermore, we did not find significant changes in expression of genes important for nonsense mediated decay (data not shown) or upregulation of this pathway. Instead, our data revealed an enrichment of an alternatively spliced *Mdm2* transcript, that was previously shown to lead to P53-stabilization and increased P53 activity (Giglio et al., 2010). Similar changes in *Mdm2* splicing were also found in the O9-1 neural crest cell line, and in patient LCLs. Hence, we postulate that this is the key event which leads to increased P53 activity in mammalian cells.

### Increased P53 activity and neural crest cell death are responsible for craniofacial malformations in the *Eftud2;Wnt1-Cre2* mutant mouse model

Genes in the P53 pathway that are upregulated in *Eftud2* mutant neural crest cells, such as *Ccng1*, *Phlda3* and *Trp53inp1* are important executors in apoptosis and proliferation. *Ccng1* encodes for Cyclin G1 and was first identified as a P53 transcriptional target (Okamoto and Beach, 1994). It is associated with a G2/M phase arrest and apoptosis induced by DNA damage in a p53-dependent manner (Kimura and Nojima, 2002). *Phlda3* encodes for Pleckstrin Homology Like Domain Family A Member 3 and is a P53-regulated repressor of Akt (Kawase et al., 2009). *Trp53inp1* encodes for Tumor protein p53-inducible nuclear protein 1, an antiproliferative and proapoptotic protein involved in cell stress response that acts as an antioxidant and plays a major role in p53-driven oxidative stress response (Cano et al., 2009). Expression of these genes are likely responsible for the death of *Eftud2* mutant neural crest cells and the profound disruption of craniofacial cartilage formation by E14.5.

The question of phenotypic variability most likely reflects the quantity and/or developmental timepoint at which these cells die. Hence, in embryos where the majority of neural crest cells die before differentiation, structures are likely to be absent, like we found for Meckel’s cartilage, while death of a lesser number of these these cells would lead to hypoplasia and/or malformed cartilages and bones. This is supported by our work with pifithrin-α which indicates that decreasing P53 activity leads to almost normal morphogenesis of the midbrain and improve the size of the pharyngeal arches. However, as many transcripts were misregulated in these mutant embryos, it remains to be determined how far development can be pushed and if a similar increase in P53 activity contributes to other craniofacial defects that are associated with mutations in splicing factors.

Our current working model is that mutations which reduce levels of EFTUD2 lead to increased exclusion of exon 3 of *MDM2.* We think that this splicing event directly or indirectly leads to activation of P53. P53 is an established stress response pathway, which provides a cell suffering from stress a chance to correct the error, in this case, a reduction in spliceosome activity. In the absence of an efficient response, the cells are then eliminated via cell death. Thus, mutant neural crest cells can be generated, migrate, colonize and differentiate in the developing craniofacial region. Determining whether MFDM clinical abnormalities would be alleviated solely by modulating P53 signaling, with the cancer risks it poses, or in combinations with other strategies to restore the other developmental networks that are surely also disrupted in these patients, will be of great importance.

## Acknowledgments

We would like to thank Dr Mitra Cowan, Associate Director of the transgenic core facility at McGill University, Goodman Cancer Center, for the micro-injections experiments. We thank the patients who agreed to participate in the study. KB and ML received support from the Care4Rare Canada Consortium funded by Genome Canada and the Ontario Genomics Institute (OGI-147), the Canadian Institutes of Health Research, Ontario Research Fund, Genome Alberta, Genome British Columbia, Genome Quebec, and Children’s Hospital of Eastern Ontario Foundation. JF received funding from NIH grant 2R15DE026611-02. This work was supported by a grant from the Canadian Institutes of Health Research (MOP#142452) given to LJAM and JM. LJAM and JM are members of the Research Centre of the McGill University Health Centre which is supported in part by FQRS. We would like to thank all members of the Jerome-Majewska lab for their unvaluable input in the writing of the manuscript.

## Author’s contribution

MCB carried out and analyzed all of the mouse experiments, drafted the manuscript and designed the figures. AD and EB performed the data analysis from the RNAseq experiments. FM carried out and analyzed all the mouse neural crest cell line experiments. RA performed IHC experiments and part of the cartilage analysis. AST did the experiments using the patients cell lines. KB, ML recruited patients involved in the study. PS supervised the analysis of experiments using patients cell lines. JF supervised the analysis of experiments using the O9-1 mouse neural crest cell line. JM supervised all bioinformatics analysis from the RNAseq. JM and LAJM devised the project. LAJM conceptualized and supervised all the mouse experiments, and wrote the manuscript. All authors provided critical feedback and helped shape the research, analysis and manuscript.

## Declaration of interests

The authors declare no competing interests.

## Materials & Methods

### Mouse lines

All procedures and experiments were performed according to the guidelines of the Canadian Council on Animal Care and approved by the Animal Care Committee of the Montreal Children’s Hospital. Wild type CD1 mice were purchased from Charles Rivers (strain code 022) and wild type C57Bl/6 mice were purchased from Jackson Laboratories (stock no 000664). The *R26R* strain (Gt(ROSA)26Sor^tm1Sor^ stock#0033O9 from Jackson’s Lab) was a kind gift from Dr Nagano. These mutant mice carry a loxP flanked DNA STOP sequence upstream preventing expression of the downstream *lacZ* gene. When crossed with a *cre* transgenic strain, the STOP sequence is removed and lacZ is expressed in cre-expressing tissues (Soriano, 1999). *Wnt1-Cre2* mice were purchased from Jackon’s laboratory (stock# 022137). These *Wnt1-Cre2* transgenic mice express Cre recombinase under the control of the mouse *Wnt1*, wingless-related MMTV integration site 1, promoter and enhancer (Lewis et al., 2013).

### Generation of *Eftud2* mouse line using CRISPR/Cas9 gene-editing system

gRNAs and microinjections for the CRISPR/Cas9 strategy used to generate the conditional mutant *Eftud2* line were described previously (Beauchamp et al., 2019).Briefly, two single stranded DNA template (ssDNA:custom-made from IDT) containing loxP sequences, EcoR1 and EcoRV flanking exon 2 of *Eftud2* were purchased. On a first round of micro-injection using gRNAs 1-4 and both ssDNA templates, one female with EcoR1/loxP insertion in the intron upstream of exon 2 was recovered. She was mated to a wild-type C57/Bl6 male and one of her male offspring with Sanger-sequenced verified insertion of the EcoR1/loxP template was used for another round of micro-injection with gRNAs 3 and 4 and the EcoRV/loxP ssDNA targeting the intron downstream of exon 2. From this second round of injections, 1 male and 1 female with EcoR1 and EcoR5 digest products were identifed. Both were mated with wild-type CD1 mice to generate G1. Intact loxP sequences were Sanger-sequenced verified in the G1 offspring of the founder male (Fig. S8). Thereafter, a G1 female carrying sequence-verified loxP sequences flanking exon 2 of *Eftud2* was backcrossed to wild-type CD1 or wild-type C57BL/6 mice for 5 generations to establish the *Eftud2* conditional mutant mouse. Embryos generated from mating of heterozygous mice from the 5^th^ generation onwards were analyzed in this study. *Eftud2 ^loxP/+^* mice were mated with each other to obtain *Eftud2 ^loxP/loxP^*.

### Generation of neural crest cell-specific deletion mutation in *Eftud2*

We first introduced the *Wnt1-Cre2* transgene in constitutive *Eftud2 ^+/-^* mice to obtain *Eftud2^+/-^; Wnt1-Cre2^tg/+^*. These mice were then mated with mice carrying conditional loxP mutations: *Eftud2 ^loxP/loxP^.* The embryos obtained (*Eftud2 ^loxP/-^; Wnt1-Cre2 ^tg/+^*) have homozygous *Eftud2* mutation in neural crest cells and heterozygous *Eftud2* mutation in all other cells. Alternatively, we also mated *Wnt1-Cre2^tg/+^* mice with *Eftud2 ^loxP/+^* to obtain *Eftud2^loxP/+^; Wnt1-Cre2^tg/+^*. These mice were then mated with mice carrying conditional loxP mutations: *Eftud2 ^loxP/loxP^.* The embryos obtained (*Eftud2 ^loxP/loxP^; Wnt1-Cre2 ^tg/+^*) have homozygous *Eftud2* mutation in neural crest cells and are wild-type in all other cells.

### Genotyping of mice

Genomic DNA was extracted from mouse tails or yolk sacs as previously described (Hou et al., 2017). For *Eftud2,* genotyping was performed using 3-primers targeting exon 2 of *Eftud2* using the following program: 30 sec 95°C, 30 sec 55°C, 30 sec 72°C for 35 cycles followed by an elongation step of 10 minutes at 72°C. As depicted in Fig. S8, the PCR condition was optimized to amplify a wild-type (180bp), a mutant (265bp) and a conditional *Eftud2* (214bp) amplicon. The following primers were used: *Eftud2* F1: atgaaccagggcagagaagt, *Eftud2* R1: tccaacagtagccaagccat, *Eftud2* R2: ccatgatgctaaaattcaaggag. For the commercially available lines, namely *Wnt1-Cre2* and *R26R*, genotyping was conducted as detailed on Jackson’s laboratory website: protocol# 29915 (*R26R*) and #25394 (*Wnt1-Cre2*).

### Collection of embryos

For embryo collection, the day of the presence of vaginal plug was considered embryonic day 0.5 (E0.5). Embryos were collected and yolk sacs were used for genomic DNA extraction for genotyping. For stages E8.5 to E10.5, the number of somites were counted under light microscope (Leica MZ6 Infinity1 stereomicroscope) at the time of dissection. Embryos were fixed in 4% paraformaldehyde at 4°C overnight (unless otherwise stated), washed in PBS and kept at 4°C.

### Cartilage preparation

To evaluate cartilage formation, embryos were stained with Alcian Blue as previously described (Rigueur and Lyons, 2014). Briefly, E14.5 embryos were fixed in Bouin’s fixing solution for 2 hours followed by several washes in 70% ethanol. Embryos were then stain in an Alcian blue staining solution (0.03% Alcian blue in 80% ethanol; 20% acetic acid) for 24-48h at 37°C. Finally, embryos were cleared in BABB (benzyl alcohol: benzyl benzoate, 1:2) solution and vizualized under light microscope (Leica MZ6 Infinity1 stereomicroscope).

### Probe production

For *Sox10* probe, the plasmid was a kind gift from Dr Trainor (Jones et al., 2008). Briefly, it was linearized with BamH1 and transcribed using T3 polymerase or linearized with EcoR1 and transcribed with T7 polymerase to generate the sense and antisense probes, respectively. Transcription of probes was done using the DIG RNA Labeling Mix (Roche). All protocols were used according to the manufacturer’s instructions.

### Preparation of embryos for *in situ* hybridization, embedding and histology

Dissected embryos were fixed in 4% paraformaldehyde overnight and dehydrated using a graded methanol series for wholemounts. Wholemount *in situs* were performed as previously described (Revil and Jerome-Majewska, 2013). For morphological analysis, E14.5 and E17.5 embryos were first decalcified in 10% EDTA for 2 weeks, then dehydrated and embedded in paraffin. Ten micrometer sections were performed on a Leica RM 2155 microtome and mounted on coated slide for further staining with hematoxylin and eosin. For cryo-embedding, fixed embryos were first cryoprotected in 30% sucrose overnight and sectioned at 10μm thickness for IF and IHC.

### Wholemount β-galactosidase staining

Embryos were fixed in 0.4% paraformaldehyde for approximately 45 minutes, washed in detergent rinse solution (0.02% Igepal, 0.01% sodium deoxycholate, 2mM MgCl_2_ in 0.1M phosphate buffer pH 7.5) 3 times 15 minutes and stain by immersion in freshly prepared 1mg/ml X-gal staining solution (0.02% Igepal, 0.01% Sodium Deoxycholate, 5mM Potassium Ferricyanide, 5mM Potassium Ferrocyanide, and 2mM MgCl_2_ in 0.1M phosphate buffer pH 7.5) overnight at 37°C in the dark. Embryos were washed in PBS and post-fix in 4% paraformaldehyde in PBS at 4°C overnight. For sectioning, embryos were embedded in Sheldon Cryomatrix (Thermoscientific) before storage in -80°C.

### Wholemount immunohistochemistry

Embryos were fixed in 0.4% PFA overnight at 4C, washed and dehydrated in increasing concentration of methanol and kept at -20C until use. Staining was performed as previously described (Jerome-Majewska et al., 2010). Briefly, after rehydration in PBS, embryos were bleached ON in methanol:DMSO:H_2_O_2_ (4:1:1) solution. They were incubated with 2H3 (DHSB 1:150) and with mouse HRP-secondary antibody (1:500) and developed with DAB. Embryos were then vizualized under light microscope (Leica MZ6 Infinity1 stereomicroscope).

### Immunofluorescence and TUNEL assay

Immunofluorescence experiments were performed according to standard protocols (Zakariyah et al., 2011). The following primary antibodies were used: Phosphohistone H3 (Ser10) (1:200 dilution, Millipore, 06–570). Alexa Fluor 568 (ThermoFisher, 1:500 dilutions) secondary antibody was used. For the quantification of apoptosis, the In Situ Cell Death Detection Kit, TMR red was used (Roche, cat# 12 156 792 910) and manufacturer’s protocol was followed. Slides were mounted withVECTASHIELD hard-set mounting medium with DAPI to visualize the nuclei. Images were captured on a Leicamicrosystem (model DM6000B) and Leica camera (model DFC 450). For quantification of fluorescence signal, particle analysis on Image J was used.

### O9-1 cell culture

O9-1 cells were cultured and maintained as described (Nguyen et al., 2018). LIF (HY-P7084), b-FGF (HY-P7004), and Mitomycin C (HY-13316) were purchased form MedChem Express. The only deviation from the protocol consisted of collecting conditioned basal O9-1 media from STO cells over the course of 5 days. All cells used for experimentation had passage numbers less than 7 and were passaged at least twice before any given experiment. Cells were cultured in 24-well plates for all experiments.

### Transfection of O9-1 cells with siRNAs

O9-1 cells were cultured to be 40% - 60% confluent on the day of transfection. *Eftud2* and negative control siRNAs (5 pmol) were transfected using the Lipofectamine® RNAiMAX transfection kit following the manufacturer’s protocol and recommendations. The cells were rinsed in cold 1X PBS and collected in 500 μl of TRIzol reagent (cat# 15596026, Thermo Fischer Scientific) 36 hours post-transfection. RNA was extracted following the manufacturer’s protocol and recommendations. For the rescue experiment, 7.5 hours after the above mentioned transfection, the cells were transfected with 1.5 μg pCAGIG or pCAGIG_Mdm2. pCAGIG was a gift from Connie Cepko (Matsuda and Cepko, 2004)(Addgene plasmid # 11159; http://n2t.net/addgene:11159; RRID: Addgene11159). To generate pCAGIG_Mdm2, the coding sequence of *Mdm2* (consisting of exons 3 through 12) was amplified with the following primers: F:GTTAGACCAAAACCATTGCTTT and R: CTAGTTGAAGTAAGTTAGCACAATCA using the Q5® High-Fidelity DNA Polymerase and Q5® Reaction Buffer and cloned into the EcorV site of pCAGIG. The cells were incubated for 36 hours after transfection with the pDNAs and RNA was extracted as described above.

### Annexin V cell death assay

O9-1 cells were cultured in 24-well plates to be 50%-60% confluent on the day of transfection. Eftud2 and negative control siRNAs were transfected using Lipofectamine® RNAiMAX as described previously. The cells were incubated in a 37C humidified incubator (5% CO2) for 24 hours post-transfection. Following the PBS wash, the cells were washed twice with 1X AnnexinV binding buffer (cat# V13246, Thermo Fischer Scientific). The AnnexinV staining solution (cat# A13201, Thermo Fisher Scientific) was added and incubated in the dark, for 15 minutes. Fluorescence was measure using a microplate reader with Ex: 495nm, ex: 519nm.

### RNA isolation for RNA sequencing and RT-qPCR

RNA extraction from samples described in Table S6 and S7 was performed using Qiagen RNeasy kit following manufacturer’s protocol. An aliquot was sent for RNA sequencing analysis. Total RNA was treated with DNAse (NEB, according to manufacturer’s protocol) and used for reverse transcription with the iScript™cDNA synthesis kit (Bio-rad, Cat. #170-8890, according to the manufacturer’s protocol). qRT-PCR was performed using the ssoAdvanced universal SYBR green supermix (Bio-Rad, cat#172-5270) on a Roche LightCycle 480 PCR machine. qPCR experiments were performed in duplicates to ensure technical replicability. Target genes were normalized with the normalization factor as calculated by geNorm software (v3.4; Ghent university hospital center for medical genetics)(Vandesompele et al., 2002). Three house-keeping genes including B2M, GAPDH, and SDHA were used for generation of the normalization factor as previously reported (Vandesompele et al., 2002). RT-PCR program included a hot start at 95 °C for 5 min, followed by 40 cycles of a denaturation step at 95 °C for 10s and an annealing/extension step at 60 °C for 30s. Primers used are listed in Table S8.

### RNA sequencing

Sequencing libraries were prepared by Genome Quebec Innovation Centre (Montreal, Canada), using the TruSeq Stranded Total RNA Sample Preparation Kit (Illumina TS-122-2301, San Diego, California, United States) by depleting ribosomal and fragmented RNA, synthesizing first and second strand cDNA, adenylating the 3′ ends and ligating adaptors, and enriching the adaptor-containing cDNA strands by PCR. The libraries were sequenced using the Illumina HiSeq 4000 PE100 sequencer, 100 nucleotide paired-end reads, generating approximately between 31 and 63 million reads sample. The sequencing reads were trimmed using CutAdapt (Martin, 2011) and mapped to the mouse reference genome (mm10) using STAR (Dobin et al., 2013) aligner (version 2.4.0e), with default parameters, and annotated using the Gencode (Mudge and Harrow, 2015) M2 (version M2, 2013) annotation. htseq-count (part of the ‘HTSeq’ (Anders et al., 2015) framework, version 0.5.4p5.) was used for expression quantification.

To perform a differential splicing analysis, we used rMATS 4.0.2 (Shen et al., 2014). To determinate if there is a difference in skipping exons between heterozygous and homozygous samples we used the Skipped Exon events reported by rMATS. The events that had a mean of inclusion junction counts (IJC) and a mean of skipped junction counts (SJC) lower than 10 in homozygous or heterozygous samples were removed from the analysis. We used the inclusion level supplied by rMATS as a difference between the average of inclusion level in heterozygous samples and the average of inclusion level in homozygous samples. We computed the number of events with an inclusion level difference more than 0.1 or less than -0.1, and a nominal p-value < 0.05 (uncorrected for multiple testing). The rationale for not using FDR correction is to obtain a large dataset enriched for alternative splicing in order to observe general tendencies, such as for example increased propensity for exon skipping or intron retention in the mutants.

To further explore intron retention events, we used IRFinder (Middleton et al., 2017) with default settings and a nominal p-value < 0.05, again motivated by the goal to detect general trends in splicing changes.

We used DESeq2 (Love et al., 2014) for differential expression analysis. Genes with a mean of read counts less or equal to 10 in homozygous and heterozygous samples were removed from the analysis. A gene set list of interest was then derived by applying filters on gene expression, such that only those with p-adj < 0.05 and FoldChange > 0 and p-adj < 0.05 and FoldChange < 0 were selected as up- and down-regulated genes.

A DAVID (version 6.8) (Huang et al., 2007) pathway analysis was performed using the list of up- and down-regulated genes. The genes used for differential expression analysis were used as background.

### Using RNA-seq data to estimate the proportion of neural crest cells

We represent the total amount of *Eftud2 <u>T</u>*ranscripts *<u>o</u>*bserved in the *<u>c</u>*ontrol samples as

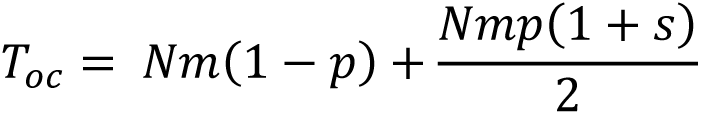

where *N* is the total number of cells, *m* is the number of transcripts produced per cell, *p* is the *<u>p</u>*roportion of neural crest cells in the head region, and *s* is the relative *<u>s</u>*tability of the *Eftud2ΔExon2* product, which is expected to introduce a frameshift and may be less stable than the complete mRNA. The first term in the equation represents transcripts obtained from normal cells (*wt*), and the second term transcripts from the neural crest fraction (which are *wt*/*Eftud2ΔExon2*). To determine the proportion of *<u>n</u>*ormal *Eftud2* transcripts, which is equivalent to the <u>P</u>ercent-<u>S</u>pliced-In (*PSI*) value for inclusion of exon 2 estimated by rMATS, we find:

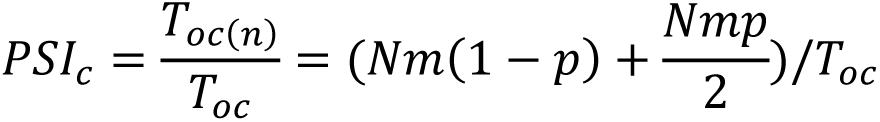

And rearranging

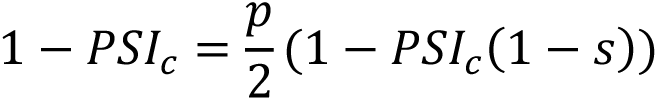

Similarly, for the *<u>m</u>*utant embryos, where the non-neural crest cells are (*wt*/*Eftud2ΔExon2*) and neural crest cells are (*Eftud2ΔExon2*/*Eftud2ΔExon2*).

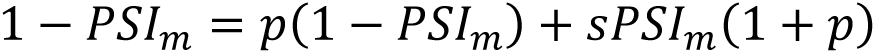

At E9.0, before the occurrence of any phenotypic defects, we assume that the proportion of neural crest cells (*p*) is the same in mutant and control embryos, and the relative stability (*s*) of the mutant *Eftud2* product is a constant parameter to be estimated. The *PSI* values in control and mutants estimated by rMATS are (*PSI*_*c*_ = 0.93; *PSI*_*m*_ = 0.51), and the above two simultaneous equations can be solved for *s* and *p.* Because of the non-linear *(sp)* term the solutions are quadratic, but only one set of solutions (*s* = 0.65; *x* = 0.21) corresponds to physiologically relevant parameters.

At E9.5, when phenotypic defects become apparent, we can use the corresponding *PSI* values (*PSI*_*c*_ = 0.84; *PSI*_*m*_ = 0.53) and the relative stability value (*s =* 0.65) determined above to estimate the proportion of neural crest cells separately in control (*p_c_*) and mutant embryos (*p_m_*) and show that, while this proportion increases in the controls (*p_c_ = 0.45*), it is further reduced in the mutants (*p_m_ = 0.15*).

### Primers used for splicing analysis

For *Mdm2* splicing analysis, cDNA was amplified with a RT-PCR program that included a hot start at 95°C for 5 min, followed by 35 cycles of a denaturation step at 95°C for 10s and an annealing step at 55°C for 30s, an extension step at 72°C for 45s with a final extension step at 72°C for 10 min, and visualized on a 2% agarose gel. The primers used are as follow: Forward (exon2-6/7): GATCACCGCGCTTCTCCTGC, Reverse (exon2-6/7): GATGTGCCAGAGTCTTGCTG (Van Alstyne et al., 2018).

### Immunohistochemistry

P53 primary antibody was used (1:250, cat#2524, Cell Signaling) and the VECTASTAIN® Universal Quick HRP Kit (PK-8800, Vector Laboratories) was used as secondary antibody and vizualized with DAB (Vector Laboratories). After rinsing with water, slides were mounted with an aqueous mounting medium and images were captured on a Leicamicrosystem (model DM6000B) and Leica camera (model DFC 450).

### LCL RNA isolation for RNA sequencing and RT-qPCR

Patient cell lines were obtained from the Care4Rare Canada Consortium with informed consent (Protocol 11/04E; Research Ethics Board of the Children’s Hospital of Eastern Ontario) (Table S3). Total RNA from WT and MFDM patient derived lymphoblastoid cell lines (LCL) was isolated using RNeasy Plus Mini Kit (Qiagen, Cat. #74134) following manufacturer’s protocol. An aliquot was sent for RNA sequencing analysis. For RT-qPCR, RNA from LCLs was reverse transcribed to cDNA using Transcriptor First Strand cDNA Synthesis Kit (Roche, Cat. # 04896866001). RT-qPCR was performed in triplicate using Fast SYBR Green Master Mix (ThermoFisher Scientific, Cat. #4385612) and StepOnePlus Real-Time PCR system (Applied Biosystems). Primers used are listed in Table S8.

### LCL RNA sequencing

Library construction was performed on total RNA (RIN ≥9.0 assessed with Agilent 6000 Nano analysis, Agilent Technologies), using pipeline for poly(A)-selected/ribo-depleted strand-specific mRNA-seq library construction from the Michael Smith Genome Sciences Centre. Biological triplicates or duplicates (Table S3) of RNA from each cell line were sequenced on a HiSeq 2500 (Illumina) using 75bp paired-end sequencing.

### Statistical analysis

Two-tailed unpaired t-test analysis was calculated using Excel and Prism Software and Chi-square test was calculated using Prism. Significant p-values are represented as *P<0.05, **P<0.01 and ***P<0.001.

All unique/stable reagents generated in this study are available from the Lead Contact with a completed Materials Transfer Agreement.

## Notes

https://www.ncbi.nlm.nih.gov/geo/query/acc.cgi?acc=GSE157356

### Competing Interest Statement

The authors have declared no competing interest.

