## Supplemental Data for "Mis-splicing of *Mdm2* leads to Increased P53-Activity and Craniofacial Defects in a MFDM *Eftud2* Mutant Mouse Model"

### Slide 1
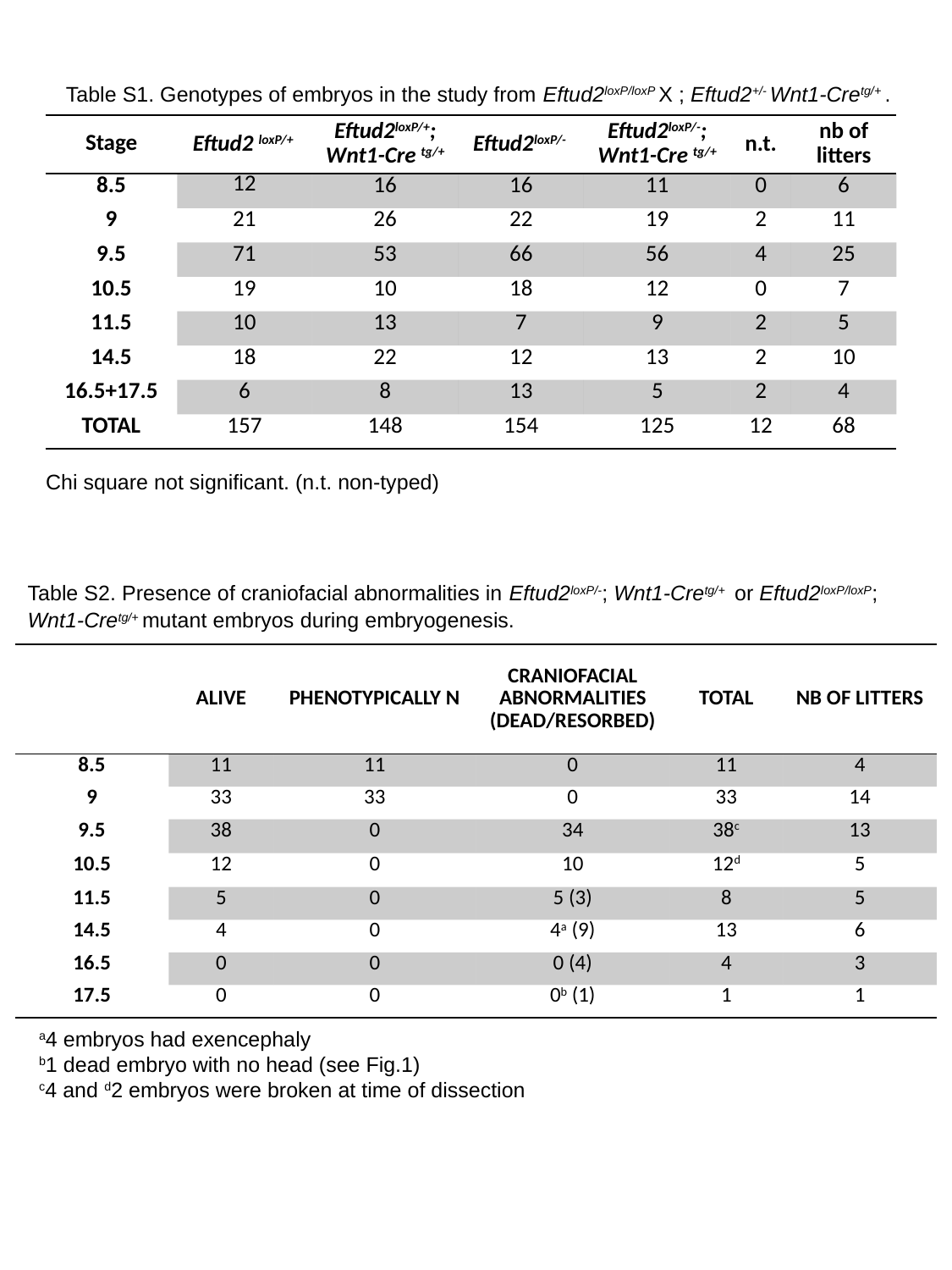

Table S1. Genotypes of embryos in the study from Eftud2^loxP/loxP^ X ; Eftud2^+/-^ Wnt1-Cre^tg/+^ .
| Stage | Eftud2 ^loxP/+^ | Eftud2^loxP/+^; Wnt1-Cre ^tg/+^ | Eftud2^loxP/-^ | Eftud2^loxP/-^; Wnt1-Cre ^tg/+^ | n.t. | nb of litters |
| --- | --- | --- | --- | --- | --- | --- |
| 8.5 | 12 | 16 | 16 | 11 | 0 | 6 |
| 9 | 21 | 26 | 22 | 19 | 2 | 11 |
| 9.5 | 71 | 53 | 66 | 56 | 4 | 25 |
| 10.5 | 19 | 10 | 18 | 12 | 0 | 7 |
| 11.5 | 10 | 13 | 7 | 9 | 2 | 5 |
| 14.5 | 18 | 22 | 12 | 13 | 2 | 10 |
| 16.5+17.5 | 6 | 8 | 13 | 5 | 2 | 4 |
| TOTAL | 157 | 148 | 154 | 125 | 12 | 68 |
Chi square not significant. (n.t. non-typed)
Table S2. Presence of craniofacial abnormalities in Eftud2^loxP/-^; Wnt1-Cre^tg/+^ or Eftud2^loxP/loxP^; Wnt1-Cre^tg/+^ mutant embryos during embryogenesis.
| | alive | phenotypically N | craniofacial abnormalities (dead/resorbed) | TOTAL | nb of litters |
| --- | --- | --- | --- | --- | --- |
| 8.5 | 11 | 11 | 0 | 11 | 4 |
| 9 | 33 | 33 | 0 | 33 | 14 |
| 9.5 | 38 | 0 | 34 | 38^c^ | 13 |
| 10.5 | 12 | 0 | 10 | 12^d^ | 5 |
| 11.5 | 5 | 0 | 5 (3) | 8 | 5 |
| 14.5 | 4 | 0 | 4^a^ (9) | 13 | 6 |
| 16.5 | 0 | 0 | 0 (4) | 4 | 3 |
| 17.5 | 0 | 0 | 0^b^ (1) | 1 | 1 |
^a^4 embryos had exencephaly
^b^1 dead embryo with no head (see Fig.1)
^c^4 and ^d^2 embryos were broken at time of dissection

### Slide 2
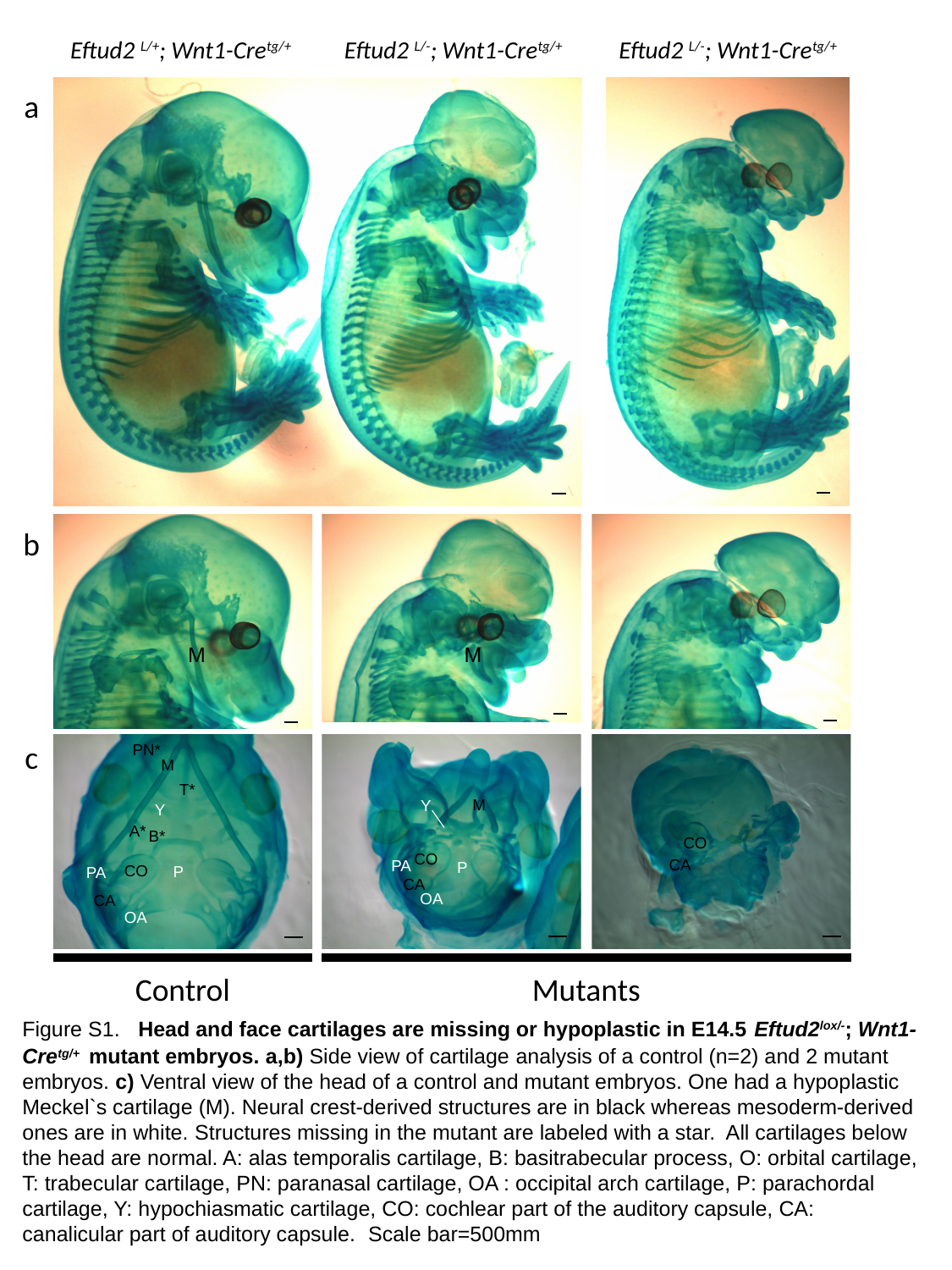

Eftud2 ^L/+^; Wnt1-Cre^tg/+^
Eftud2 ^L/-^; Wnt1-Cre^tg/+^
Eftud2 ^L/-^; Wnt1-Cre^tg/+^
a
b
M
M
c
PN*
M
T*
M
Y
Y
A*
B*
CO
CO
CA
PA
P
CO
P
PA
CA
OA
CA
OA
Control
Mutants
Figure S1. Head and face cartilages are missing or hypoplastic in E14.5 Eftud2^lox/-^; Wnt1-Cre^tg/+^ mutant embryos. a,b) Side view of cartilage analysis of a control (n=2) and 2 mutant embryos. c) Ventral view of the head of a control and mutant embryos. One had a hypoplastic Meckel`s cartilage (M). Neural crest-derived structures are in black whereas mesoderm-derived ones are in white. Structures missing in the mutant are labeled with a star. All cartilages below the head are normal. A: alas temporalis cartilage, B: basitrabecular process, O: orbital cartilage, T: trabecular cartilage, PN: paranasal cartilage, OA : occipital arch cartilage, P: parachordal cartilage, Y: hypochiasmatic cartilage, CO: cochlear part of the auditory capsule, CA: canalicular part of auditory capsule. Scale bar=500mm

### Slide 3
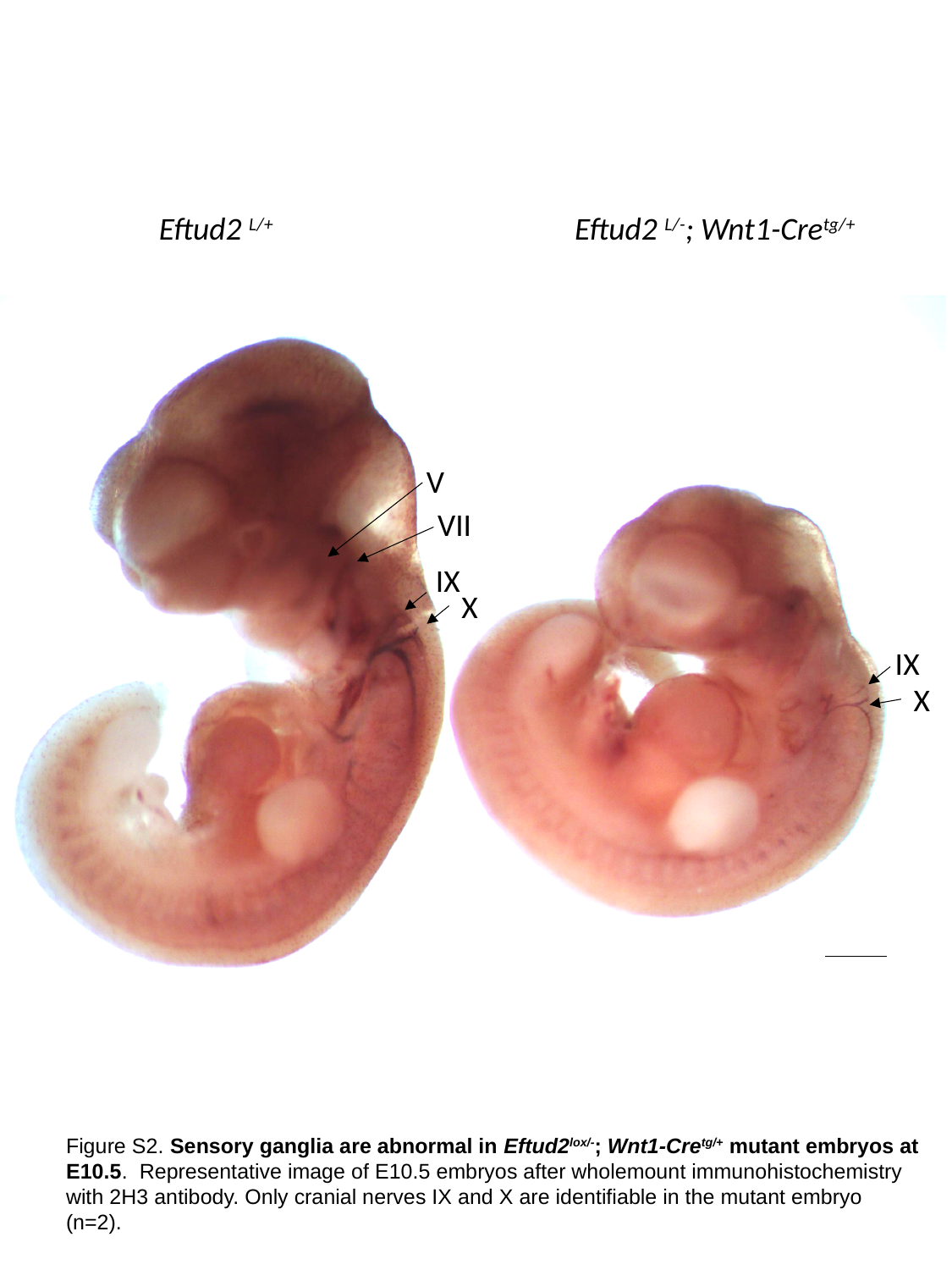

Eftud2 ^L/+^
Eftud2 ^L/-^; Wnt1-Cre^tg/+^
V
VII
IX
X
IX
X
Figure S2. Sensory ganglia are abnormal in Eftud2^lox/-^; Wnt1-Cre^tg/+^ mutant embryos at E10.5. Representative image of E10.5 embryos after wholemount immunohistochemistry with 2H3 antibody. Only cranial nerves IX and X are identifiable in the mutant embryo (n=2).

### Slide 4
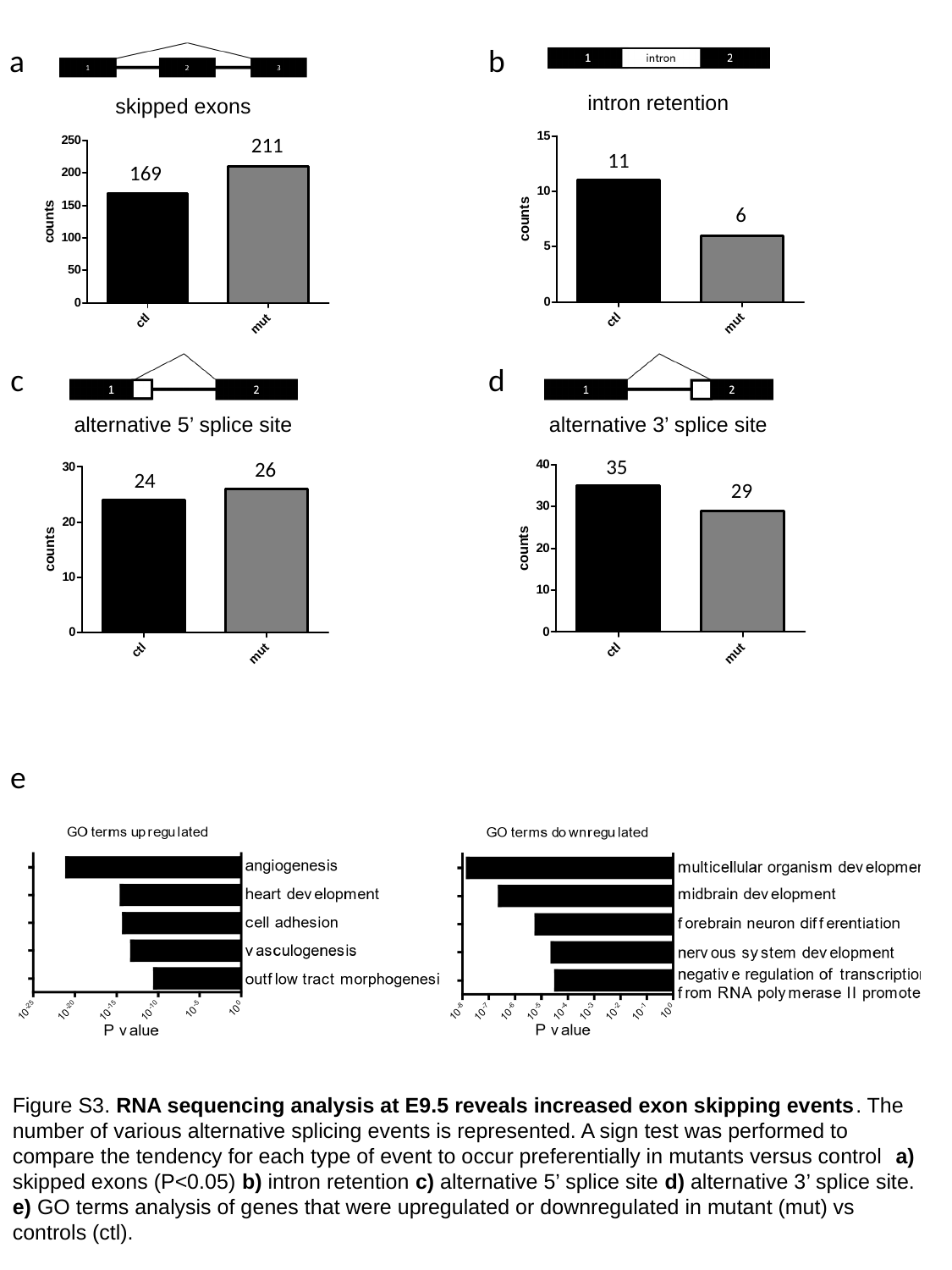

a
b
skipped exons
intron retention
11
6
211
169
c
d
alternative 5’ splice site
alternative 3’ splice site
35
29
26
24
e
Figure S3. RNA sequencing analysis at E9.5 reveals increased exon skipping events. The number of various alternative splicing events is represented. A sign test was performed to compare the tendency for each type of event to occur preferentially in mutants versus control a) skipped exons (P<0.05) b) intron retention c) alternative 5’ splice site d) alternative 3’ splice site. e) GO terms analysis of genes that were upregulated or downregulated in mutant (mut) vs controls (ctl).

### Slide 5
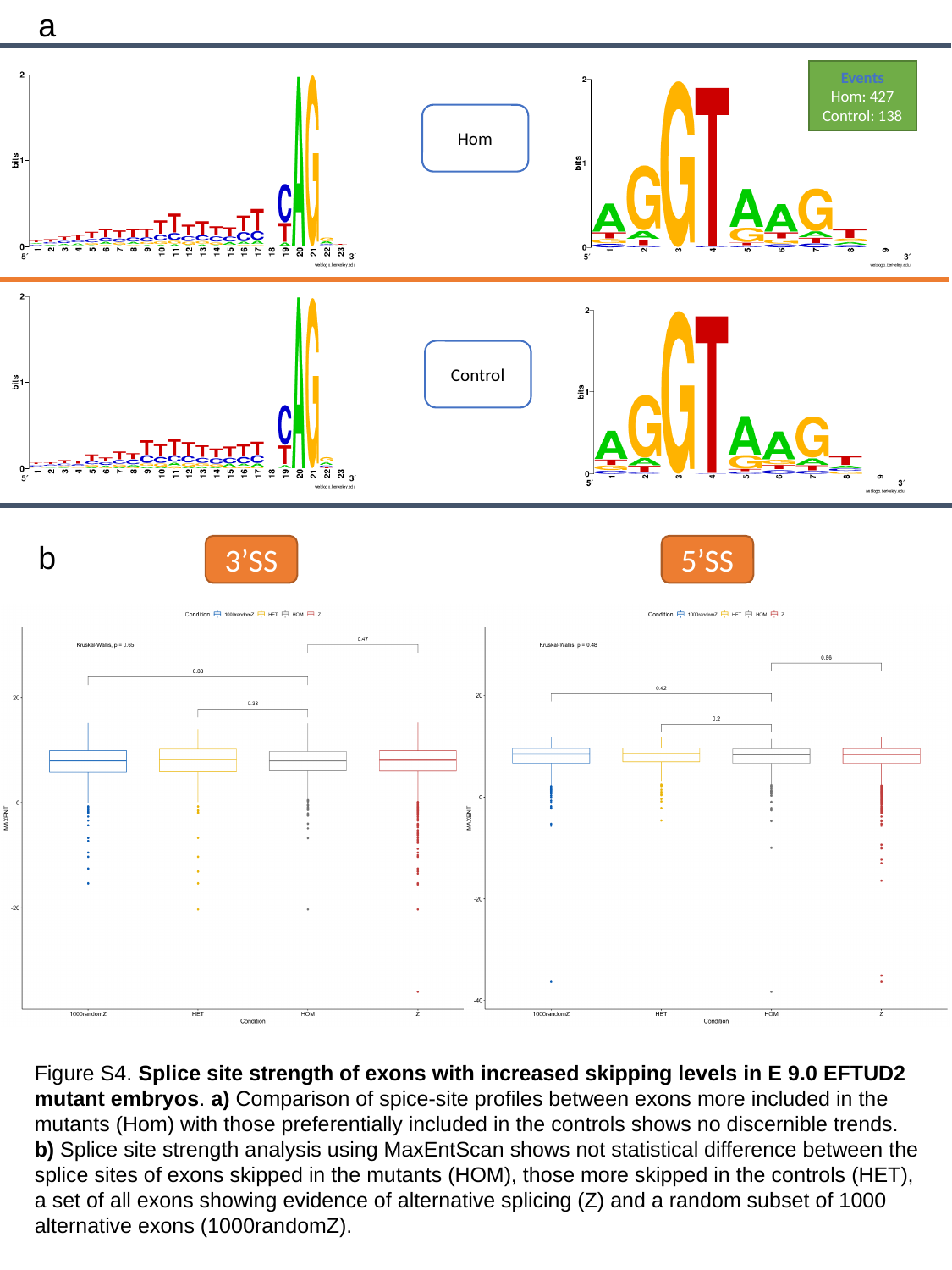

a
5’SS
Events
Hom: 427
Control: 138
Hom
Control
b
3’SS
5’SS
Figure S4. Splice site strength of exons with increased skipping levels in E 9.0 EFTUD2 mutant embryos. a) Comparison of spice-site profiles between exons more included in the mutants (Hom) with those preferentially included in the controls shows no discernible trends. b) Splice site strength analysis using MaxEntScan shows not statistical difference between the splice sites of exons skipped in the mutants (HOM), those more skipped in the controls (HET), a set of all exons showing evidence of alternative splicing (Z) and a random subset of 1000 alternative exons (1000randomZ).

### Slide 6
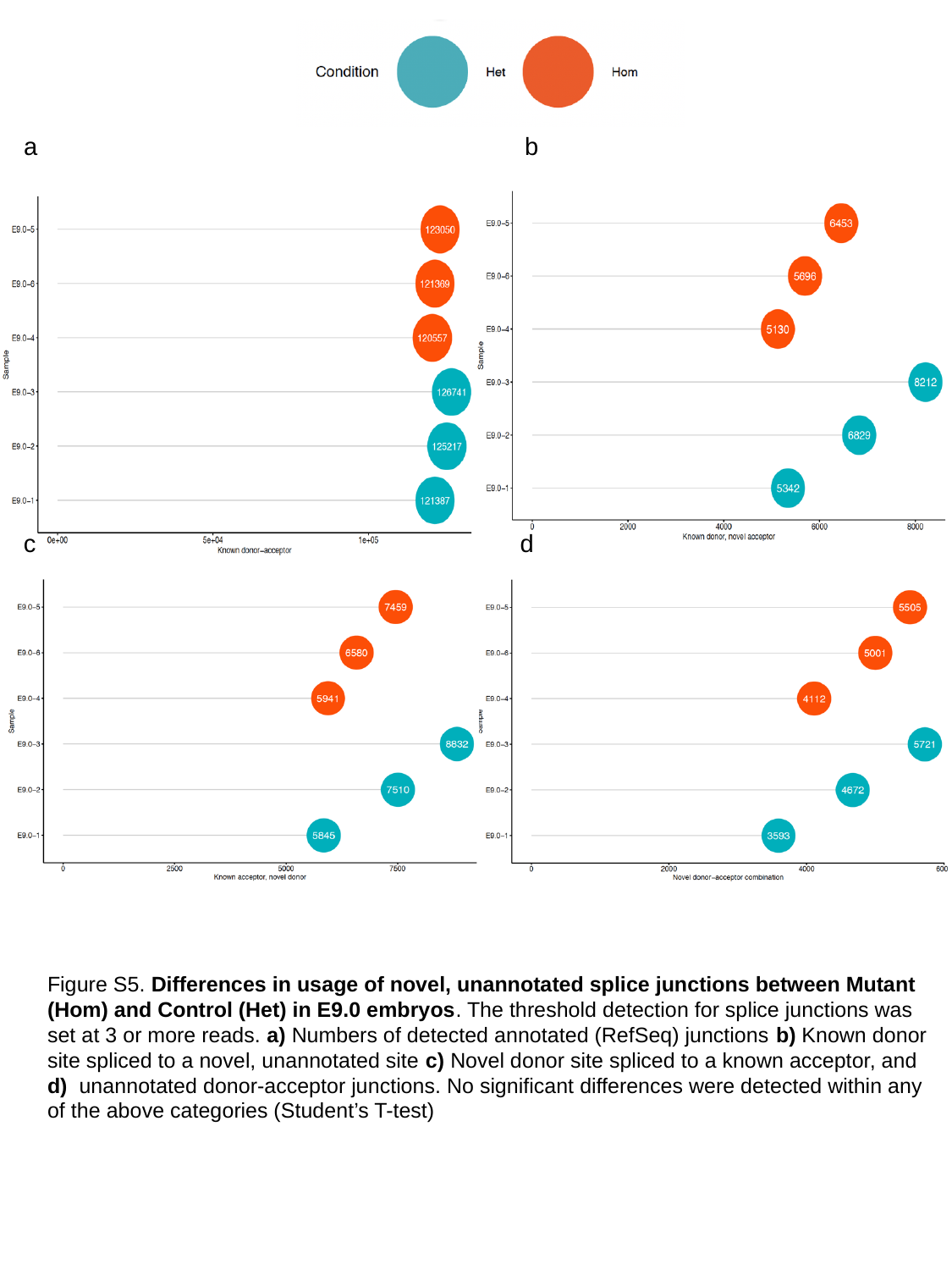

b
a
c
d
Figure S5. Differences in usage of novel, unannotated splice junctions between Mutant (Hom) and Control (Het) in E9.0 embryos. The threshold detection for splice junctions was set at 3 or more reads. a) Numbers of detected annotated (RefSeq) junctions b) Known donor site spliced to a novel, unannotated site c) Novel donor site spliced to a known acceptor, and d) unannotated donor-acceptor junctions. No significant differences were detected within any of the above categories (Student’s T-test)

### Slide 7
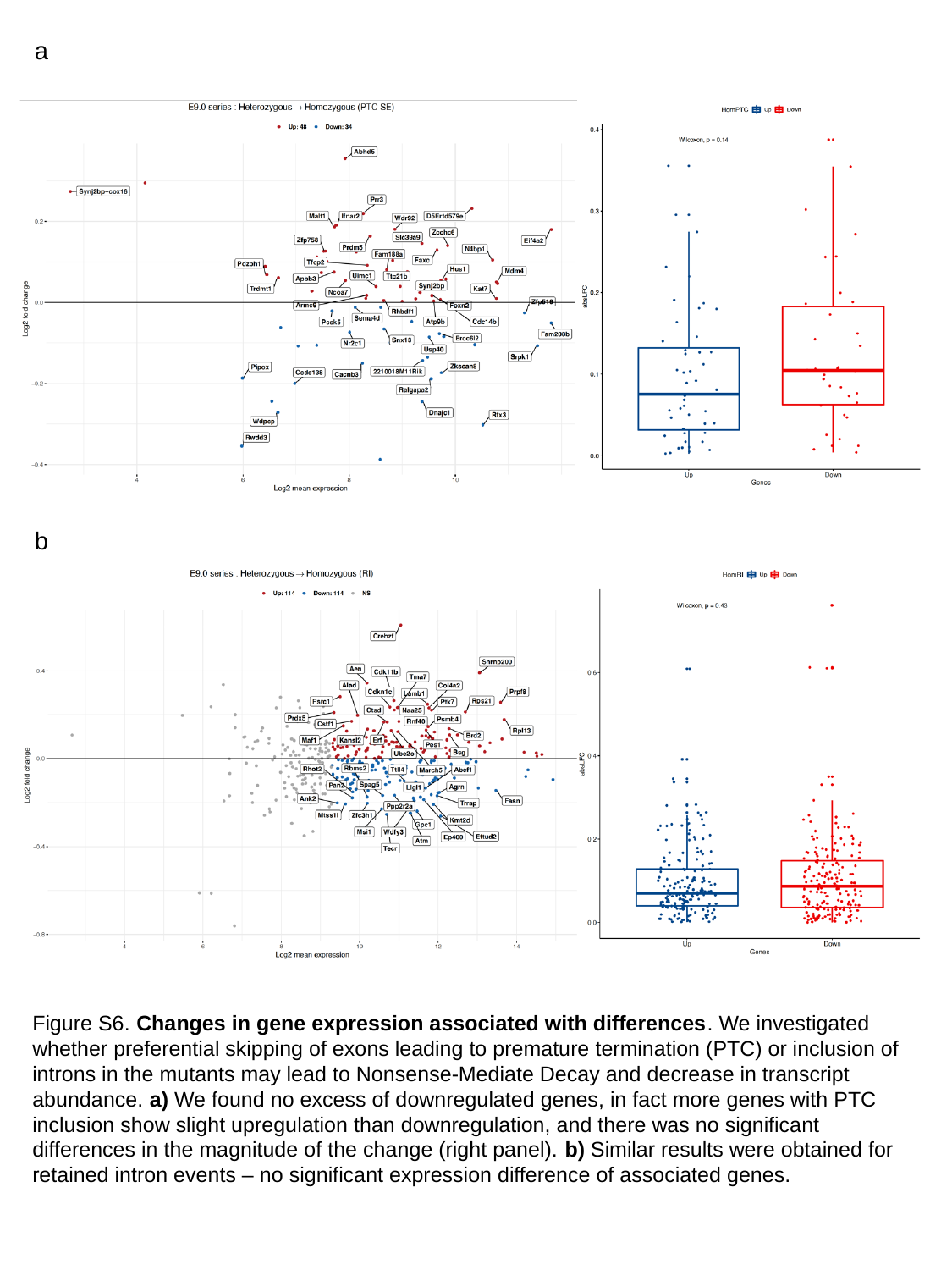

a
b
Figure S6. Changes in gene expression associated with differences. We investigated whether preferential skipping of exons leading to premature termination (PTC) or inclusion of introns in the mutants may lead to Nonsense-Mediate Decay and decrease in transcript abundance. a) We found no excess of downregulated genes, in fact more genes with PTC inclusion show slight upregulation than downregulation, and there was no significant differences in the magnitude of the change (right panel). b) Similar results were obtained for retained intron events – no significant expression difference of associated genes.

### Slide 8
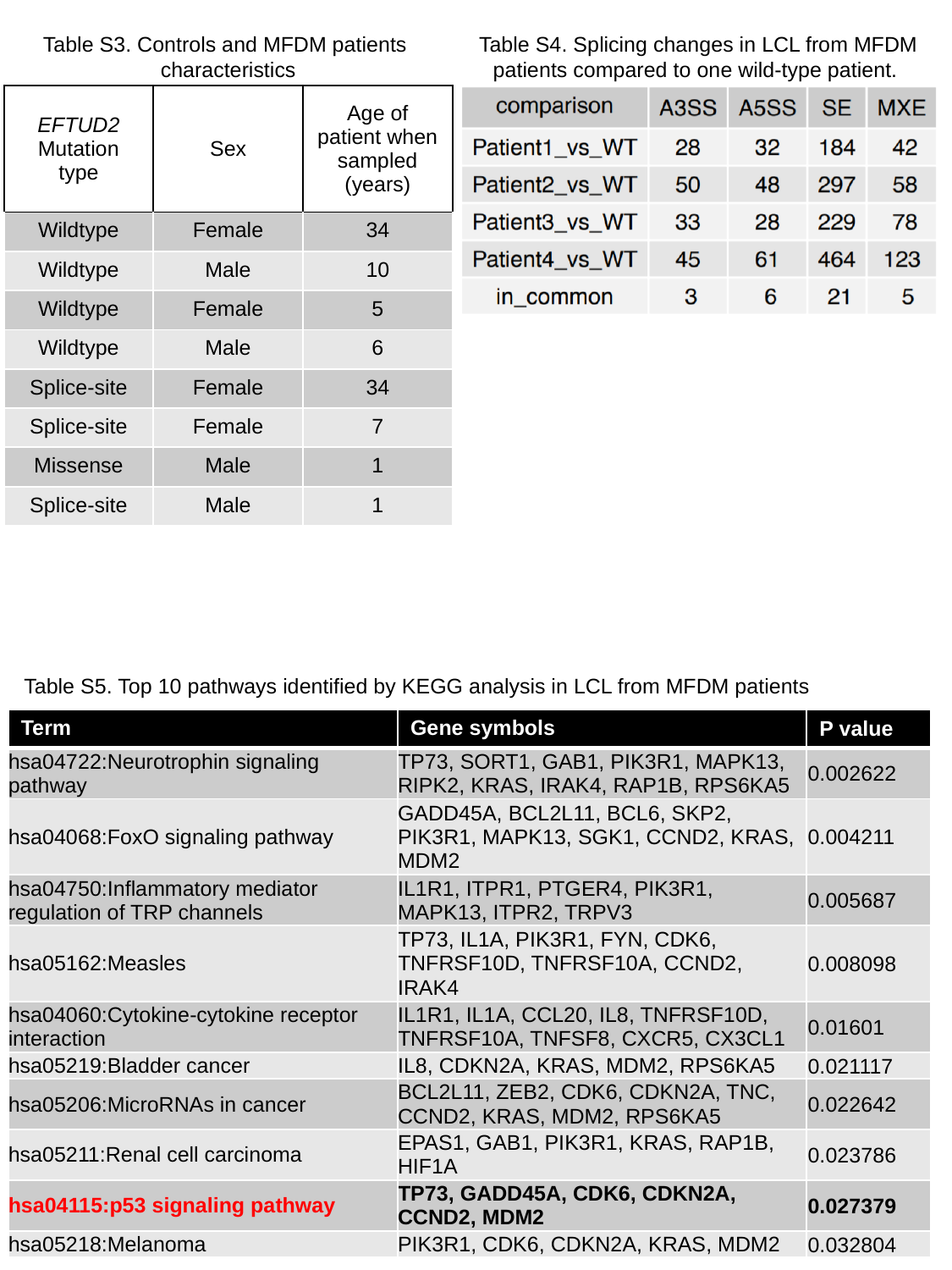

Table S3. Controls and MFDM patients
characteristics
Table S4. Splicing changes in LCL from MFDM patients compared to one wild-type patient.
| EFTUD2 Mutation type | Sex | Age of patient when sampled (years) |
| --- | --- | --- |
| Wildtype | Female | 34 |
| Wildtype | Male | 10 |
| Wildtype | Female | 5 |
| Wildtype | Male | 6 |
| Splice-site | Female | 34 |
| Splice-site | Female | 7 |
| Missense | Male | 1 |
| Splice-site | Male | 1 |
Table S5. Top 10 pathways identified by KEGG analysis in LCL from MFDM patients
| Term | Gene symbols | P value |
| --- | --- | --- |
| hsa04722:Neurotrophin signaling pathway | TP73, SORT1, GAB1, PIK3R1, MAPK13, RIPK2, KRAS, IRAK4, RAP1B, RPS6KA5 | 0.002622 |
| hsa04068:FoxO signaling pathway | GADD45A, BCL2L11, BCL6, SKP2, PIK3R1, MAPK13, SGK1, CCND2, KRAS, MDM2 | 0.004211 |
| hsa04750:Inflammatory mediator regulation of TRP channels | IL1R1, ITPR1, PTGER4, PIK3R1, MAPK13, ITPR2, TRPV3 | 0.005687 |
| hsa05162:Measles | TP73, IL1A, PIK3R1, FYN, CDK6, TNFRSF10D, TNFRSF10A, CCND2, IRAK4 | 0.008098 |
| hsa04060:Cytokine-cytokine receptor interaction | IL1R1, IL1A, CCL20, IL8, TNFRSF10D, TNFRSF10A, TNFSF8, CXCR5, CX3CL1 | 0.01601 |
| hsa05219:Bladder cancer | IL8, CDKN2A, KRAS, MDM2, RPS6KA5 | 0.021117 |
| hsa05206:MicroRNAs in cancer | BCL2L11, ZEB2, CDK6, CDKN2A, TNC, CCND2, KRAS, MDM2, RPS6KA5 | 0.022642 |
| hsa05211:Renal cell carcinoma | EPAS1, GAB1, PIK3R1, KRAS, RAP1B, HIF1A | 0.023786 |
| hsa04115:p53 signaling pathway | TP73, GADD45A, CDK6, CDKN2A, CCND2, MDM2 | 0.027379 |
| hsa05218:Melanoma | PIK3R1, CDK6, CDKN2A, KRAS, MDM2 | 0.032804 |

### Slide 9
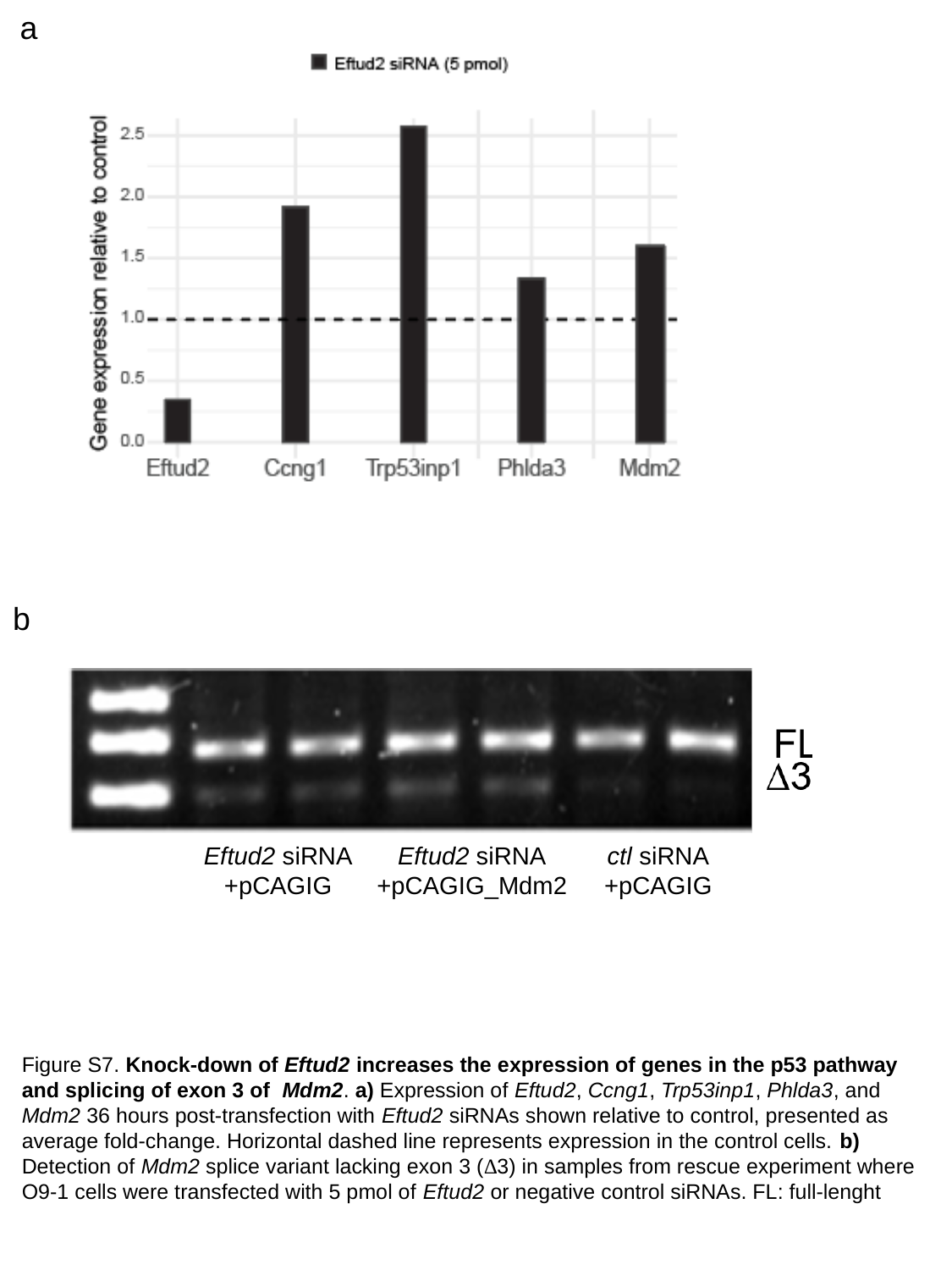

a
b
Eftud2 siRNA
+pCAGIG
Eftud2 siRNA
+pCAGIG_Mdm2
ctl siRNA
+pCAGIG
Figure S7. Knock-down of Eftud2 increases the expression of genes in the p53 pathway and splicing of exon 3 of Mdm2. a) Expression of Eftud2, Ccng1, Trp53inp1, Phlda3, and Mdm2 36 hours post-transfection with Eftud2 siRNAs shown relative to control, presented as average fold-change. Horizontal dashed line represents expression in the control cells. b) Detection of Mdm2 splice variant lacking exon 3 (∆3) in samples from rescue experiment where O9-1 cells were transfected with 5 pmol of Eftud2 or negative control siRNAs. FL: full-lenght

### Slide 10
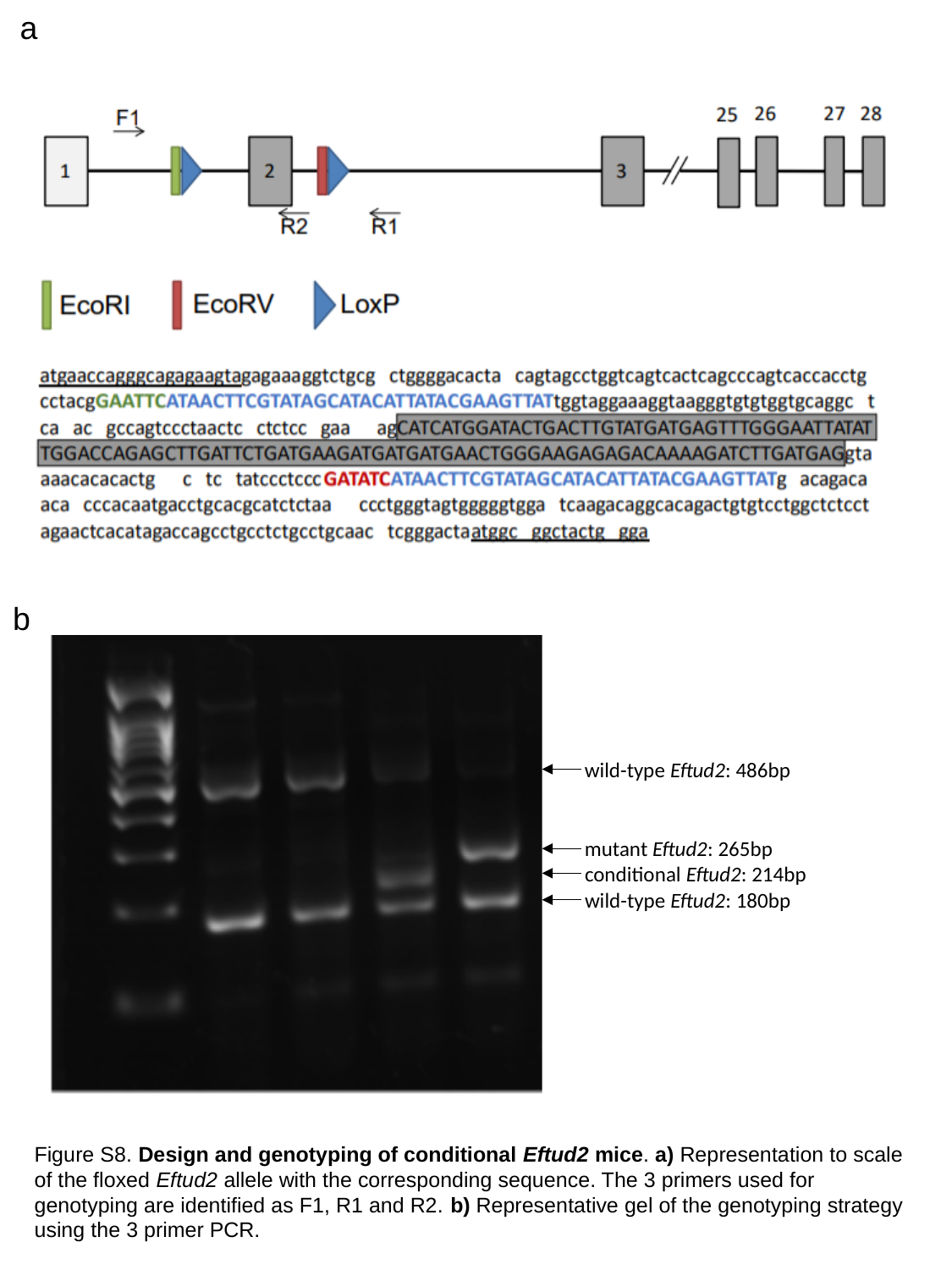

a
b
wild-type Eftud2: 486bp
mutant Eftud2: 265bp
conditional Eftud2: 214bp
wild-type Eftud2: 180bp
Figure S8. Design and genotyping of conditional Eftud2 mice. a) Representation to scale of the floxed Eftud2 allele with the corresponding sequence. The 3 primers used for genotyping are identified as F1, R1 and R2. b) Representative gel of the genotyping strategy using the 3 primer PCR.

### Slide 11
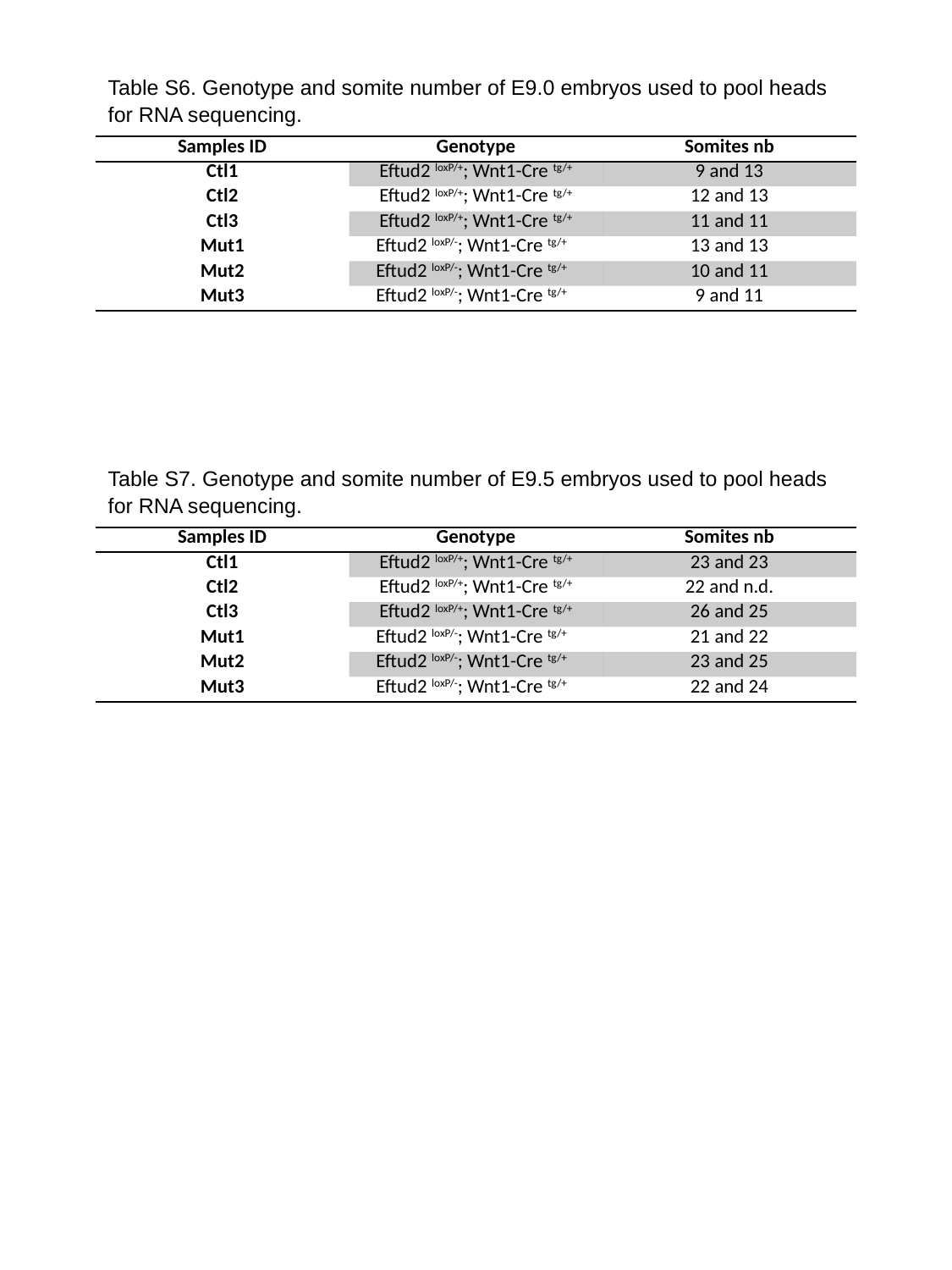

Table S6. Genotype and somite number of E9.0 embryos used to pool heads for RNA sequencing.
| Samples ID | Genotype | Somites nb |
| --- | --- | --- |
| Ctl1 | Eftud2 ^loxP/+^; Wnt1-Cre ^tg/+^ | 9 and 13 |
| Ctl2 | Eftud2 ^loxP/+^; Wnt1-Cre ^tg/+^ | 12 and 13 |
| Ctl3 | Eftud2 ^loxP/+^; Wnt1-Cre ^tg/+^ | 11 and 11 |
| Mut1 | Eftud2 ^loxP/-^; Wnt1-Cre ^tg/+^ | 13 and 13 |
| Mut2 | Eftud2 ^loxP/-^; Wnt1-Cre ^tg/+^ | 10 and 11 |
| Mut3 | Eftud2 ^loxP/-^; Wnt1-Cre ^tg/+^ | 9 and 11 |
Table S7. Genotype and somite number of E9.5 embryos used to pool heads for RNA sequencing.
| Samples ID | Genotype | Somites nb |
| --- | --- | --- |
| Ctl1 | Eftud2 ^loxP/+^; Wnt1-Cre ^tg/+^ | 23 and 23 |
| Ctl2 | Eftud2 ^loxP/+^; Wnt1-Cre ^tg/+^ | 22 and n.d. |
| Ctl3 | Eftud2 ^loxP/+^; Wnt1-Cre ^tg/+^ | 26 and 25 |
| Mut1 | Eftud2 ^loxP/-^; Wnt1-Cre ^tg/+^ | 21 and 22 |
| Mut2 | Eftud2 ^loxP/-^; Wnt1-Cre ^tg/+^ | 23 and 25 |
| Mut3 | Eftud2 ^loxP/-^; Wnt1-Cre ^tg/+^ | 22 and 24 |

### Slide 12
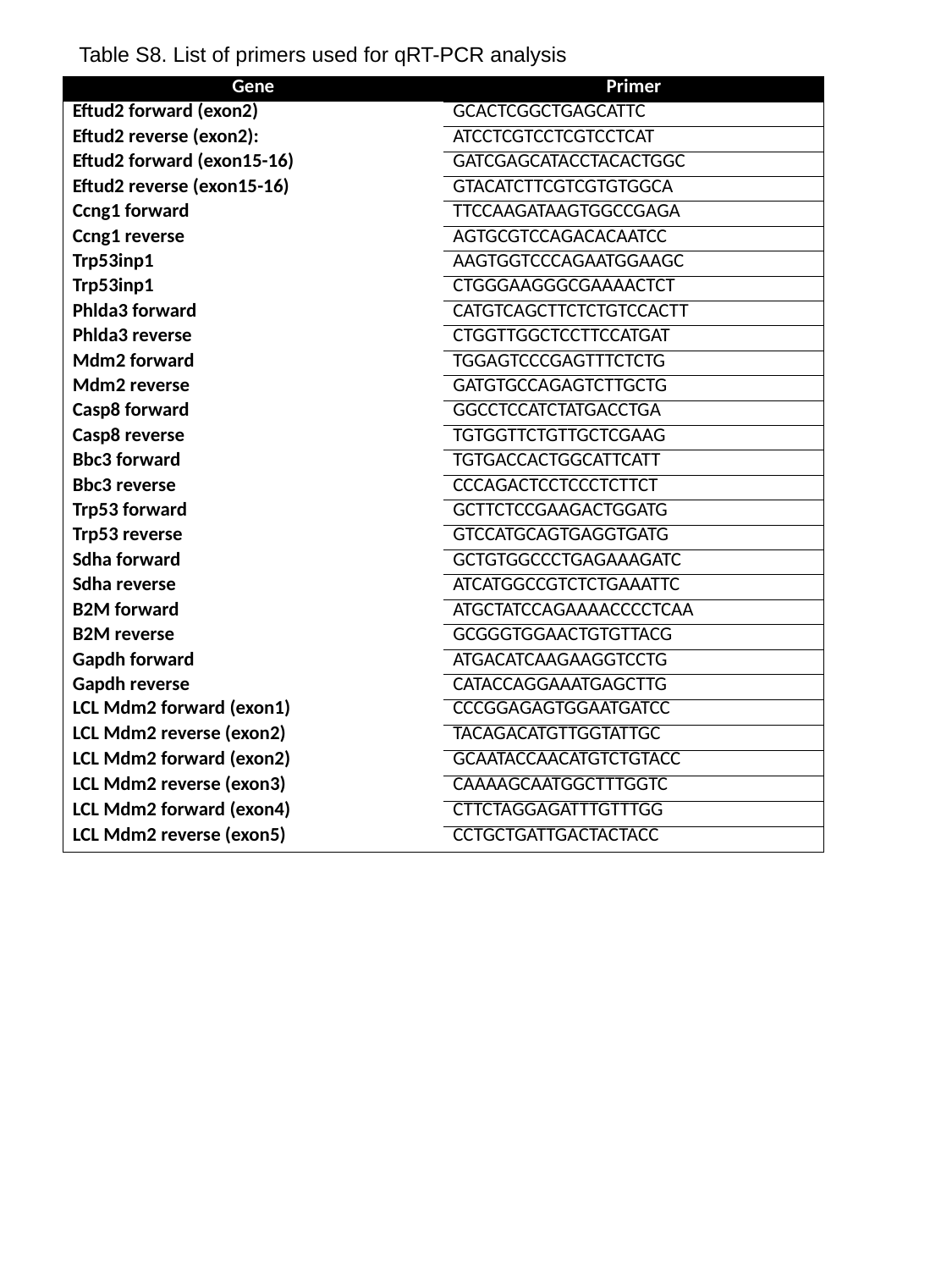

Table S8. List of primers used for qRT-PCR analysis
| Gene | Primer |
| --- | --- |
| Eftud2 forward (exon2) | gcactcggctgagcattc |
| Eftud2 reverse (exon2): | atcctcgtcctcgtcctcat |
| Eftud2 forward (exon15-16) | GATCGAGCATACCTACACTGGC |
| Eftud2 reverse (exon15-16) | GTACATCTTCGTCGTGTGGCA |
| Ccng1 forward | TTCCAAGATAAGTGGCCGAGA |
| Ccng1 reverse | AGTGCGTCCAGACACAATCC |
| Trp53inp1 | AAGTGGTCCCAGAATGGAAGC |
| Trp53inp1 | CTGGGAAGGGCGAAAACTCT |
| Phlda3 forward | CATGTCAGCTTCTCTGTCCACTT |
| Phlda3 reverse | CTGGTTGGCTCCTTCCATGAT |
| Mdm2 forward | TGGAGTCCCGAGTTTCTCTG |
| Mdm2 reverse | GATGTGCCAGAGTCTTGCTG |
| Casp8 forward | ggcctccatctatgacctga |
| Casp8 reverse | tgtggttctgttgctcgaag |
| Bbc3 forward | tgtgaccactggcattcatt |
| Bbc3 reverse | cccagactcctccctcttct |
| Trp53 forward | gcttctccgaagactggatg |
| Trp53 reverse | gtccatgcagtgaggtgatg |
| Sdha forward | GCTGTGGCCCTGAGAAAGATC |
| Sdha reverse | ATCATGGCCGTCTCTGAAATTC |
| B2M forward | ATGCTATCCAGAAAACCCCTCAA |
| B2M reverse | GCGGGTGGAACTGTGTTACG |
| Gapdh forward | ATGACATCAAGAAGGTCCTG |
| Gapdh reverse | CATACCAGGAAATGAGCTTG |
| LCL Mdm2 forward (exon1) | CCCGGAGAGTGGAATGATCC |
| LCL Mdm2 reverse (exon2) | TACAGACATGTTGGTATTGC |
| LCL Mdm2 forward (exon2) | GCAATACCAACATGTCTGTACC |
| LCL Mdm2 reverse (exon3) | CAAAAGCAATGGCTTTGGTC |
| LCL Mdm2 forward (exon4) | CTTCTAGGAGATTTGTTTGG |
| LCL Mdm2 reverse (exon5) | CCTGCTGATTGACTACTACC |
